# Systematic Interrogation Uncovers Targetable Translational Control Elements within the CFTR 5’ UTR

**DOI:** 10.64898/2026.10.16.759746

**Authors:** Ben T. Pockrass, Alice M. Youle, Rachel O. Niederer

**Author notes:** Address correspondence to R.O.N. 4560 MSRB2 1150 W Medical Center Dr Ann Arbor, MI 48109.

## Abstract

The 5’ untranslated region (5’ UTR) of mRNAs often contain features that regulate ribosome recruitment and translation. These regulatory features represent potential targets to modulate translation of therapeutically relevant genes, such as cystic fibrosis transmembrane conductance regulator (CFTR), the gene responsible for Cystic Fibrosis. However, the CFTR 5’ UTR regulatory landscape is largely unknown. To comprehensively identify regulatory elements in CFTR 5’ UTR that could be targeted to modulate its translation, we measured ribosome recruitment to a pool of systematically mutated CFTR 5’ UTRs using Direct Analysis of Ribosome Targeting (DART). This assay allowed us to test the effect of putative 5’ UTR regulatory features as well as discover novel regulatory elements in an unbiased manner. Using this data, we validated a known repressive uAUG in a fetal-lung specific CFTR 5’ UTR isoform and identified a stimulatory secondary structure in the canonical CFTR 5’ UTR. Using a large set of targeted mutations, we also discovered stimulatory (CFTR 5’ UTR Associated Stimulator of Translation, CAST) and repressive (CFTR 5’ UTR Associated Repressor of Translation, CART) sequence features in the CFTR 5’ UTR that regulate translation. Further, we found that CART is present in many human 5’ UTRs, enriched for membrane proteins and ion transporters, and acts broadly as a translational repressor, suggesting this motif may regulate translation of not only CFTR but related genes. Overall, our strategy successfully identified regulatory features within the CFTR 5’ UTR which represent promising therapeutic targets.

**Graphical Abstract:** 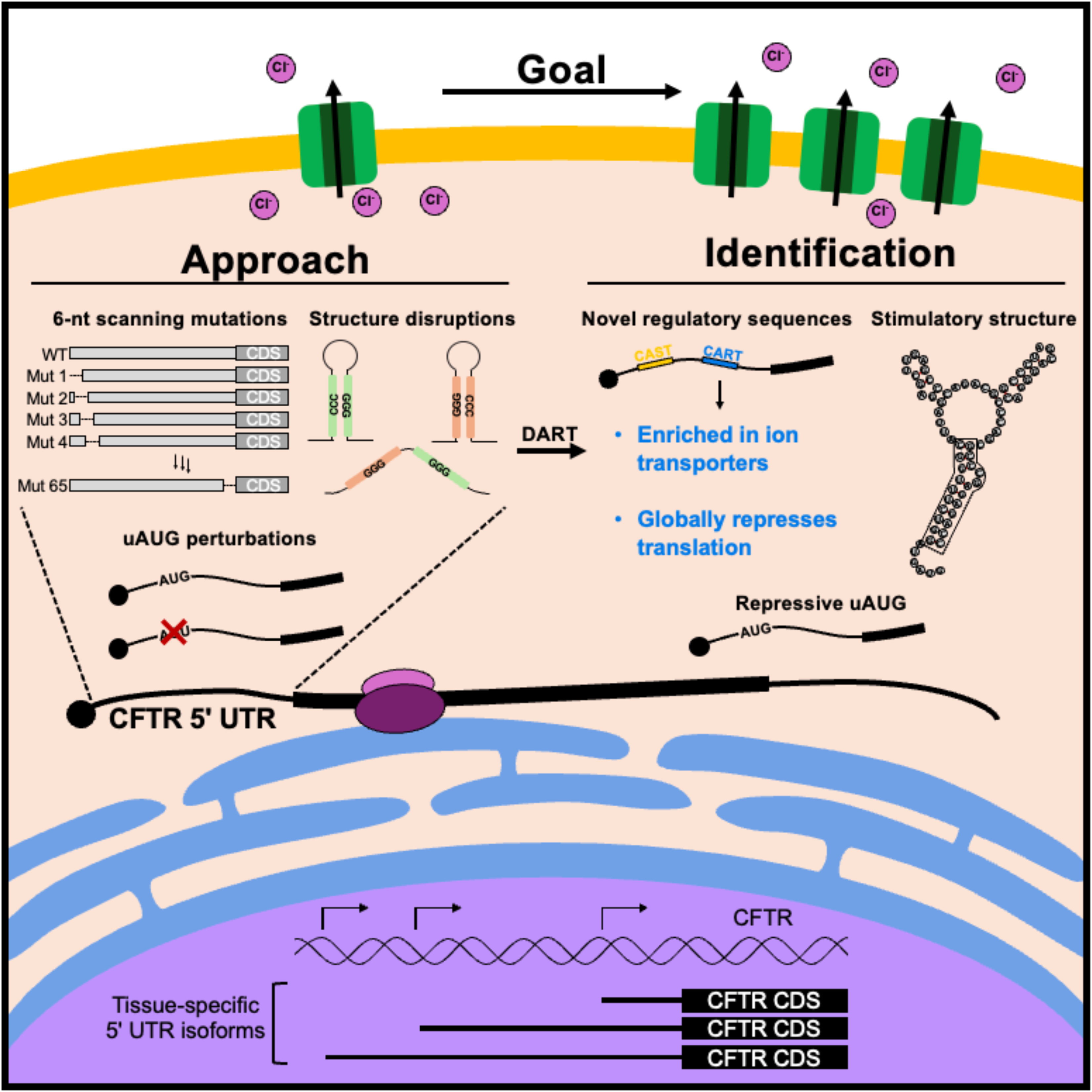

## Introduction

Precise manipulation of protein levels is a longstanding need therapeutically, and targeting mRNA translation specifically has shown promise in treating several disorders^1–12^. The 5’ untranslated region (5’ UTR) of an mRNA often contains regulatory elements^13–15^ such as secondary structures^16,17^, RNA binding protein (RBP) binding sites^18–20^, and upstream start codons (uAUG)^21–24^ that can mediate thousand fold differences in translational output^25^. We hypothesize that these features can be targeted to modulate translation of therapeutically relevant genes, but most 5’ UTRs remain largely uncharacterized. Here, we aim to uncover new regulatory features within the Cystic Fibrosis Transmembrane Conductance Regulator (CFTR) 5’ UTR that can be targeted to increase its expression (Fig. 1A). Critically, how elements in the CFTR 5’ UTR regulate its translation has not been systematically evaluated.

**Figure 1.**
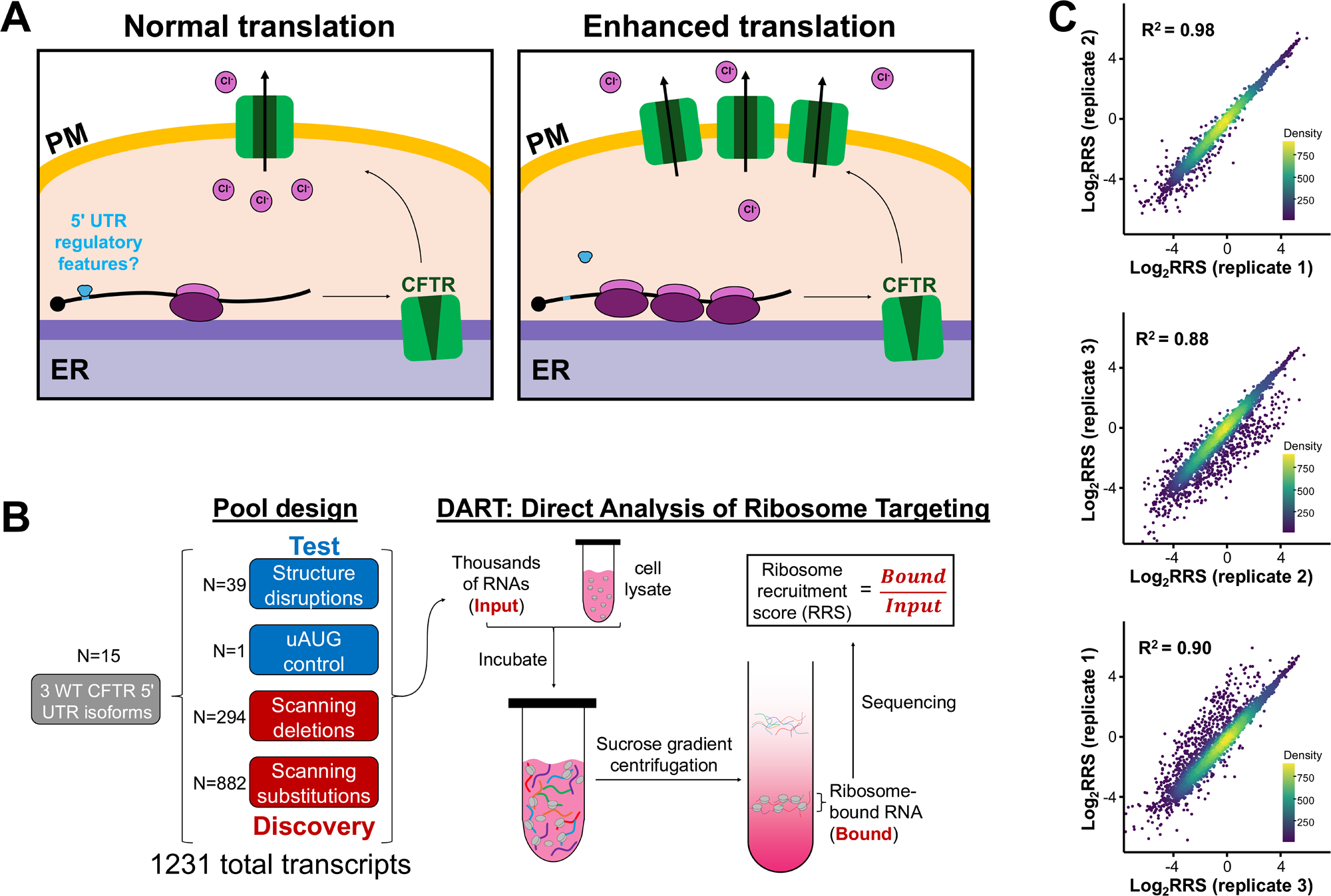
A strategy to find elements in the CFTR 5’ UTR that regulate translation. **A.** Increasing CFTR protein levels by enhancing its translation is a potential therapeutic strategy for many CF-causing variants. Translation of CFTR could be modulated by targeting features in its 5’ UTR, but these features are currently unknown. ER: endoplasmic reticulum. PM: plasma membrane. **B.** An overview of pool contents used in Direct Analysis of Ribosome Targeting (DART). Putative regulatory elements in the CFTR 5’ UTR were perturbed to evaluate their effect on ribosome recruitment (Test), and a large set of targeted mutations were introduced to the CFTR 5’ UTR to identify regions that contain regulatory elements that cannot be predicted by sequence alone (Discovery). **C.** DART was performed on a pool of mutated CFTR 5’ UTRs in triplicate. Correlations between three biological replicates are shown.

Cystic fibrosis (CF) is a genetic disease caused by defects to CFTR, an ATP-gated chloride channel expressed in various tissues including the lungs, pancreas, and intestines^26^. CFTR biogenesis is complex: it requires targeting to and cotranslational folding at the endoplasmic reticulum (ER), post translational modification, and trafficking to the plasma membrane to ultimately carry out its function^27^. Not surprisingly, there are hundreds of mutations to CFTR that cause CF, which are divided into six classes based on how the mutant affects CFTR biogenesis and/or function^28,29^. Despite different molecular effects across mutational classes, most classes also exhibit reduced protein levels. Notably, a recent study found that over 90% of exonic CFTR variants result in lower protein levels than WT^30^. Low protein levels are an obstacle even for highly effective modulator treatments, which can only act on protein that is present in the cell^30^. Therefore, strategies to increase CFTR protein levels are expected to be therapeutically beneficial across classes. Given that enhancing translation of CFTR specifically has been shown to increase functional expression^31–34^ we reasoned that approaches targeting the CFTR 5’ UTR could represent a promising, mutation-agnostic strategy.

CFTR has three main 5’ UTR isoforms which are generated by tissue-specific alternative transcription start sites^26,35^; a 70-nt isoform (CFTR-70) expressed in the lung, intestines, and pancreas, a 107-nt isoform (CFTR-107) expressed exclusively in the intestines, and a 132-nt isoform (CFTR-132) exclusively expressed in fetal lung^26^. CFTR-132 is the most studied isoform and is known to contain a repressive uAUG as well as a repressive secondary structure, with this structure also repressing translation of CFTR-70^36^. Targeting of the uAUG with antisense oligonucleotides (ASOs) in CFTR-132 has been shown to improve CFTR expression for multiple mutants^37^. Despite this, the most widely expressed and clinically relevant CFTR 5’ UTR isoform (CFTR-70) is comparatively uncharacterized.

We reasoned that a comprehensive understanding of CFTR translation regulation by its 5’ UTR would reveal features that could be targeted to increase its translation. Here, we perform Direct Analysis of Ribosome Targeting (DART) on a pool of 1216 mutated CFTR 5’ UTRs to comprehensively identify features regulating its translation. Our data confirm the previously characterized repressive uAUG in CFTR-132 as well as identify a stimulatory secondary structure in CFTR-70. Using a series of targeted mutations across the entire length of the CFTR 5’ UTRs, we also identified novel stimulatory and repressive sequences that regulate translation of CFTR-70. Finally, we show that this repressive sequence is enriched in human 5’ UTRs encoding ion transporters, and that it is associated with low ribosome recruitment and translation in other 5’ UTRs. Overall, we have successfully identified functional regulatory elements in the CFTR 5’ UTR that are attractive therapeutic targets and established a robust pipeline to discover targetable features in any mRNA of interest.

## Results

### A strategy to find regulatory features in the CFTR 5’ UTR

While CFTR expression is regulated at many stages, the full repertoire of translational control elements within the 5’ UTR has not been uncovered. To comprehensively identify features in the CFTR 5’ UTR that regulate translation, we performed Direct Analysis of Ribosome Targeting (DART)^25^, which measures ribosome recruitment to thousands of RNAs in parallel, on a pool of systematically mutated CFTR 5’ UTRs. We reasoned that disrupting functional 5’ UTR regulatory elements would result in significant differences in ribosome recruitment, which is a proxy for the translational capacity of an mRNA^25^. Systematic variations were introduced into three annotated 5’ UTR isoforms to make the DART pool (Fig. 1B, left). The pool included 40 mutations to test the effect of established translational control elements, such as secondary structures and upstream AUGs (uAUGs), and 1176 systematic disruptions across the isoforms to discover regulatory elements in an unbiased manner. Together, these comprehensively interrogate translation regulatory features within the CFTR 5’ UTR.

The RNA pool was incubated in cytoplasmic extract derived from human bronchial epithelial cells (16HBE14o-, referred to herein as HBE) followed by separation on a sucrose gradient to purify ribosome-bound RNAs (Fig. 1B, right). Ribosome-bound RNAs were sequenced and their abundance compared to the input RNA to calculate a ribosome recruitment score (RRS) for each transcript in the pool. RRS varied up to 500-fold between variants and showed high reproducibility between independent replicates (R^2^ > 0.88, Fig. 1C) enabling investigation of the regulatory landscape of the CFTR 5’ UTR.

### Tissue-specific CFTR 5’ UTR isoforms drive different levels of translation

CFTR is expressed in multiple tissues including the lung, pancreas, and intestines at tissue-specific transcription start sites^26,35^ (Fig. 2A). These alternative TSSs produce 5’ UTR isoforms that are effectively 5’ extensions of each other and vary in length from 70-nt to 132-nt. CFTR-70 is expressed in lung, pancreatic, and intestinal tissue, whereas CFTR-107 and CFTR-132 isoforms are specific to intestinal and fetal lung tissue, respectively. To determine whether these 5’ UTR isoforms can mediate levels of translation, we first compared their RRS’s (Fig. 2B). We find that the isoforms differ in their ribosome recruitment ability, with CFTR-132 showing the highest values. To determine how protein production compares between the isoforms, we generated firefly luciferase reporters containing each of the 3 CFTR 5’ UTR isoforms and measured their translational output in an *in vitro* translation assay (Fig. 2C) using HBE cell lysate. Translation from CFTR-70 was the highest, with CFTR-132 and CFTR-107 producing about half as much protein.

**Figure 2.**
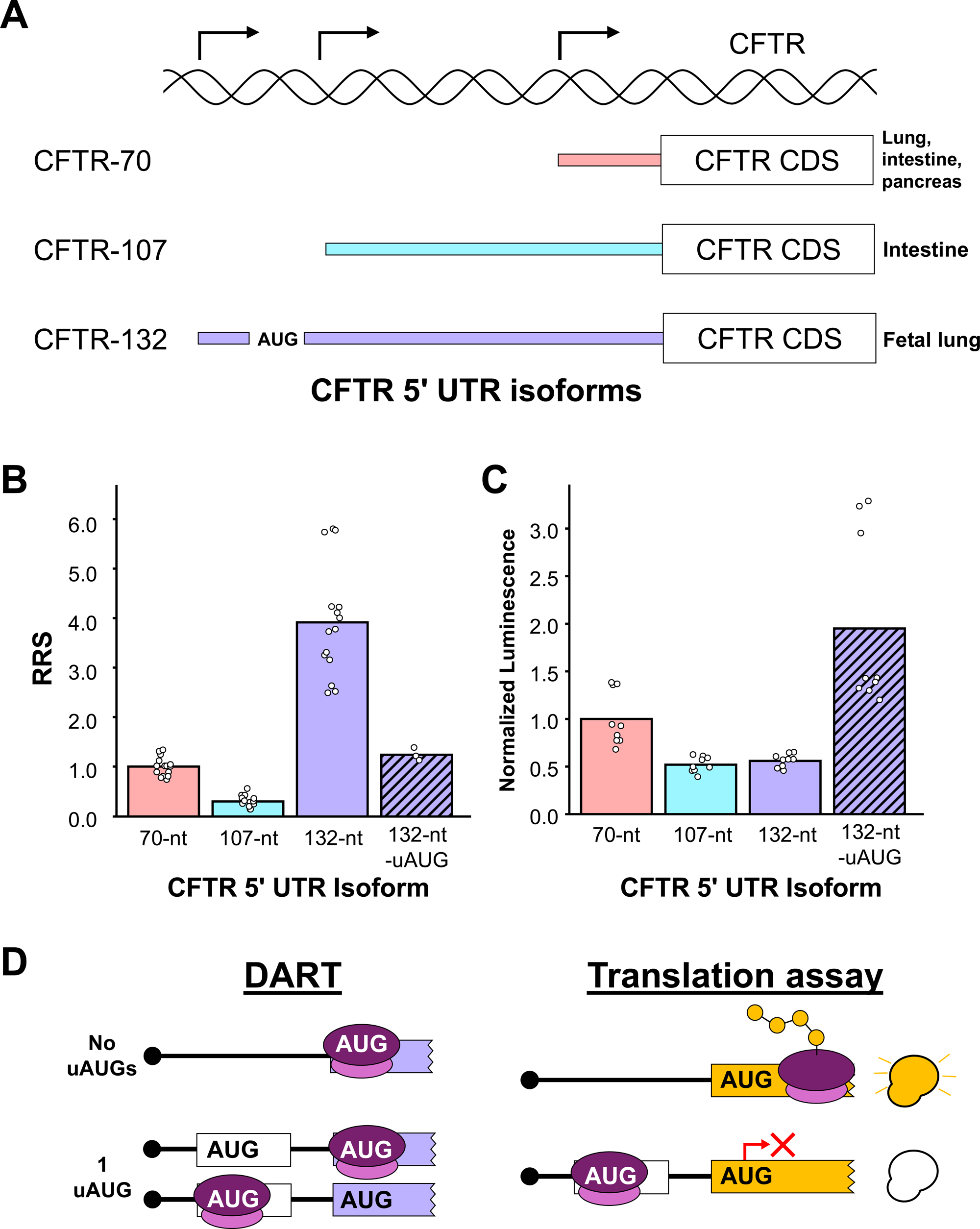
CFTR has multiple 5’ UTR isoforms which vary in their ribosome recruitment and translational output. **A.** Three tissue-specific CFTR 5’ UTR isoforms generated from alternative transcription start sites were tested for their ability to recruit ribosomes by DART. **B.** Ribosome recruitment scores (RRS) were calculated for each tested CFTR 5’ UTR isoform. Bars are averages among three biological replicates for five individually barcoded transcripts in the pool, except for the – uAUG mutant which was only included in the pool once. Data are normalized to CFTR-70. The assay was performed using HBE lysate. **C.** *In vitro* translation assays were performed on each CFTR 5’ UTR isoform tested in DART. Bars are averages among nine biological replicates. Data are normalized to CFTR-70. The assay was performed in HBE lysate. **D.** DART detects both uAUGs and AUGs, leading to relatively high RRS (left) even if an uAUG represses translation at the main AUG in a translation assay (right).

Notably, CFTR-132 is the only isoform known to contain a repressive uAUG and exhibits a relatively high RRS and low translational output. We reasoned that functional, out-of-frame uAUGs increase the likelihood of a given 5’ UTR becoming ribosome-bound in DART while reducing usage of the main AUG in a translation assay (Fig. 2D). Indeed, mutating the uAUG to AGT in CFTR-132 decreases its RRS (Fig. 2B) and increases its translational output (Fig. 2C). These data support that tissue-specific CFTR 5’ UTRs vary in their ribosome-recruitment and translational compacities, and that DART can identify functional uAUGs.

### The CFTR 5’ UTR contains a stimulatory secondary structure

Each CFTR 5’ UTR isoform is predicted to form a unique secondary structure (Fig. 3A, left, Supp. Fig. 1A, B). To test the importance of these putative structures, we made structure disrupting mutations to each individual stem loop (Fig. 3A, right, Supp. table 1). For each predicted stem, we included three mutants: a 5’ and 3’ mutant which both disrupt the stem loop by making substitutions to each side of the stem, respectively, and a compensatory mutant which incorporates both 5’ and 3’ mutations such that the predicted structure is restored. If the structure is regulating ribosome recruitment independent of sequence, we expect both 5’ and 3’ mutants to have the same effect on RRS, while the compensatory mutant should recapitulate WT RRS levels. In structure disrupting mutants to the CFTR-70 stem loop 1, we see this effect: mutating both the 5’ and 3’ ends of the stem decreases RRS, and the compensatory mutant partially rescues RRS (Fig. 3B). These data suggest that base pairing interactions within stem 1 enhance ribosome recruitment and potentially translation. Structure disrupting mutations to stems 2 and 3 have variable effects, suggesting either the predicted structures do not form or do not regulate ribosome recruitment (Fig. 3C, D).

**Figure 3.**
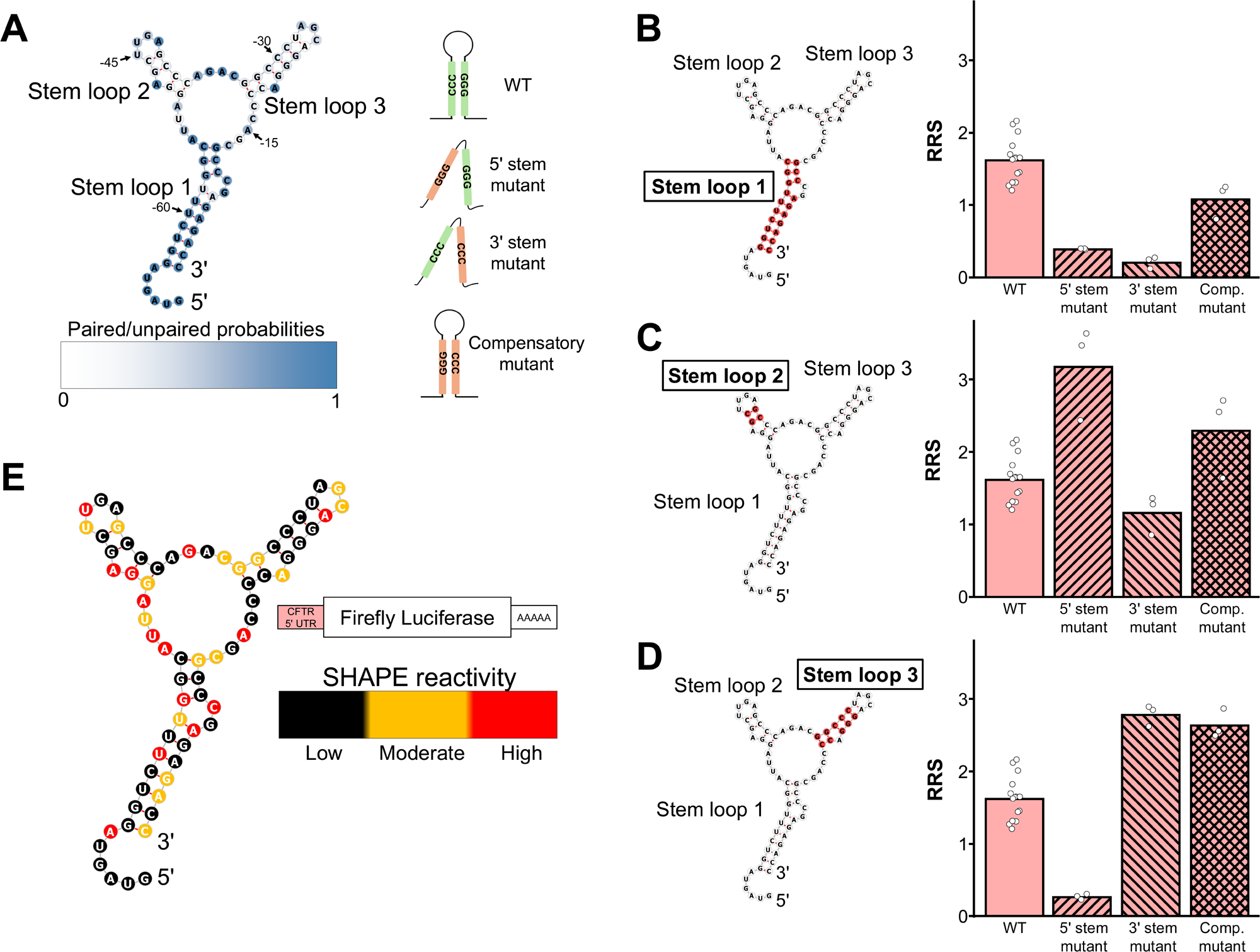
DART detects functional 5’ UTR structures. **A.** Minimum free energy RNA secondary structure was predicted using the RNAfold web server for the CFTR-70 5’ UTR and used to make targeted mutations to disrupt each predicted stem loop (see supplementary table 1 for mutants). Nucleotides are labeled according to their position with respect to the first nucleotide of the start codon. Secondary structures were visualized using forna. Each predicted stem structure was disrupted in the DART pool by making three mutants: 5’ and 3’ mutants that make substitutions to the 5’ and 3’ sequence of the stem, respectively, and a compensatory mutant that incorporates substitutions from both mutants such that a stem should still form. **B-D.** Ribosome recruitment scores (RRS) for structure disruptions were compared to the WT for stem 1 **(B)**, stem 2 **(C)**, and stem 3 **(D)** in the CFTR-70 5’ UTR. Bars represent averages of three biological replicates (dots), except the WT which is an average of three biological replicates for five individually barcoded transcripts in the pool. Nucleotides highlighted in red indicate the mutated sequence. The assay was performed in HBE lysate. **E.** SHAPE-MaP was performed on a luciferase reporter containing the CFTR-70 5’ UTR. SHAPE reactivity is indicated in the scale and mapped onto the minimum free energy prediction of the CFTR-70 5’ UTR.

To experimentally determine the structure adopted by the CFTR 5’ UTR we performed SHAPE-MaP^38^ on CFTR-70 (Fig. 3E, Supp. Fig. 2). Interestingly, we observe low reactivity throughout stem 3, suggesting this structure does form even though it does not regulate ribosome recruitment (Fig. 3D). Both stems 1 and 2 show areas of reactivity within the predicted pairing regions, suggesting either CFTR-70 adopts alternative structures in these regions or that the structures are highly dynamic.

### Substitutions have highly variable impacts on RRS

To identify 5’ UTR features regulating translation of CFTR in an unbiased manner, we generated a set of overlapping 6-nt mutations across the entire length of each CFTR 5’ UTR isoform (Fig. 4A, left). We took a two-pronged approach, either deleting the 6-nt segment or substituting it with a random 6-nt sequence. For the substitution series, three randomly generated 6-nt sequences were substituted at every 6-nt window. Scanning mutations were offset by 1-nt, such that consecutive mutations contain 5-nt overlap. We aimed to identify regions of the CFTR 5’ UTR that increase, decrease, or have no effect on ribosome recruitment by comparing the RRS of each mutant to the WT (Fig. 4A, right). To identify these features, we calculated the log_2_ of the fold change in RRS between each mutant and the WT (log_2_(RRS_mut_/RRS_WT_)), hereafter referred to as the relative RRS.

**Figure 4.**
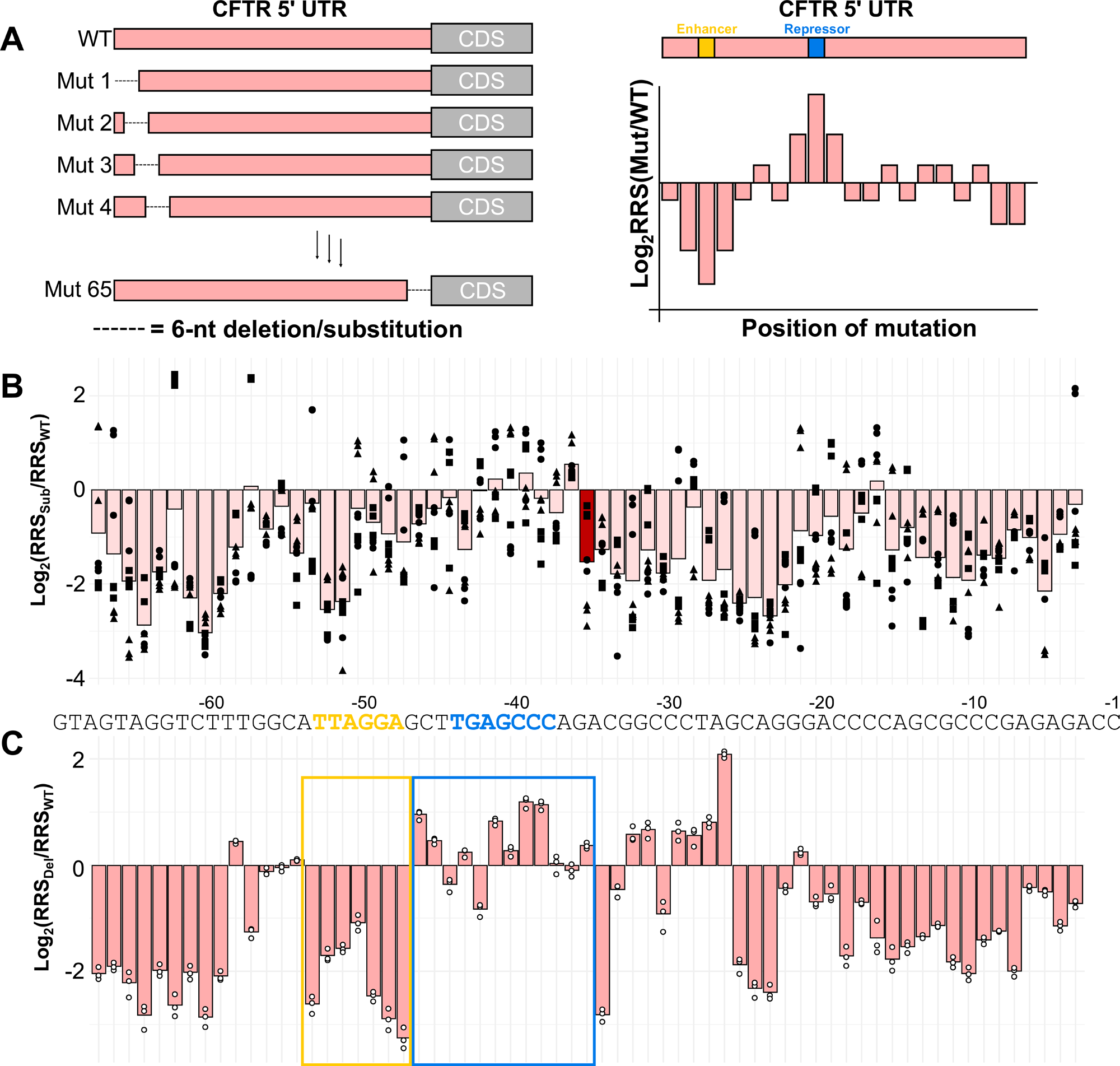
Systematic mutations across the CFTR-70 5’ UTR reveal regions that regulate translation. **A.** 6-nt scanning deletions or substitutions were introduced and shifted 1-nt at a time down the entire length of the CFTR-70 5’ UTR. Ribosome recruitment was measured to these mutants using DART. **B.** The log_2_ fold change (log_2_FC) of the RRS between each 6-nt substitution and the WT was calculated. For each 6-nt window, three different random 6-nt substitutions were made, and all were measured by DART in three biological replicates. Bars represent the average log_2_FC for all replicates, and circles, squares, and triangles distinguish the three different sequences substituted at a given position. Bars are centered on the 6-nt that are substituted (the WT CFTR-70 5’ UTR sequence shown at the bottom). The dark red bar corresponds to positions -38 to -33, which is referenced in the text. **C.** The log_2_FC of the RRS between each 6-nt deletion and the WT was calculated. Each bar represents the average log_2_FC for a given deletion for three biological replicates (dots). Bars are centered on the 6-nt that are deleted (the WT CFTR-70 5’ UTR sequence shown on top). The assay was performed in HBE lysate.

We first compared the deletion or substitution approach to determine which strategy was most interpretable. Deletion of sequences is conceptually straightforward, however it also changes the length of the UTR. Conversely, substitutions maintain a consistent length but due to the introduction of new sequence may add new translational control elements. We found that the random sequence substituted at a given position often caused large variation in the RRS and were overall difficult to interpret (Fig. 5B). For example, the relative RRS at nucleotide positions -38 to -33 are widely distributed but clearly cluster by the identity of the substituted sequence (Fig. 4B, dark red bar). Therefore, we concluded that random substitution was more indicative of gained effects from the introduced sequences rather than reporting on the disrupted sequence. Consequently, we focused our downstream analyses on the scanning deletion series.

**Figure 5.**
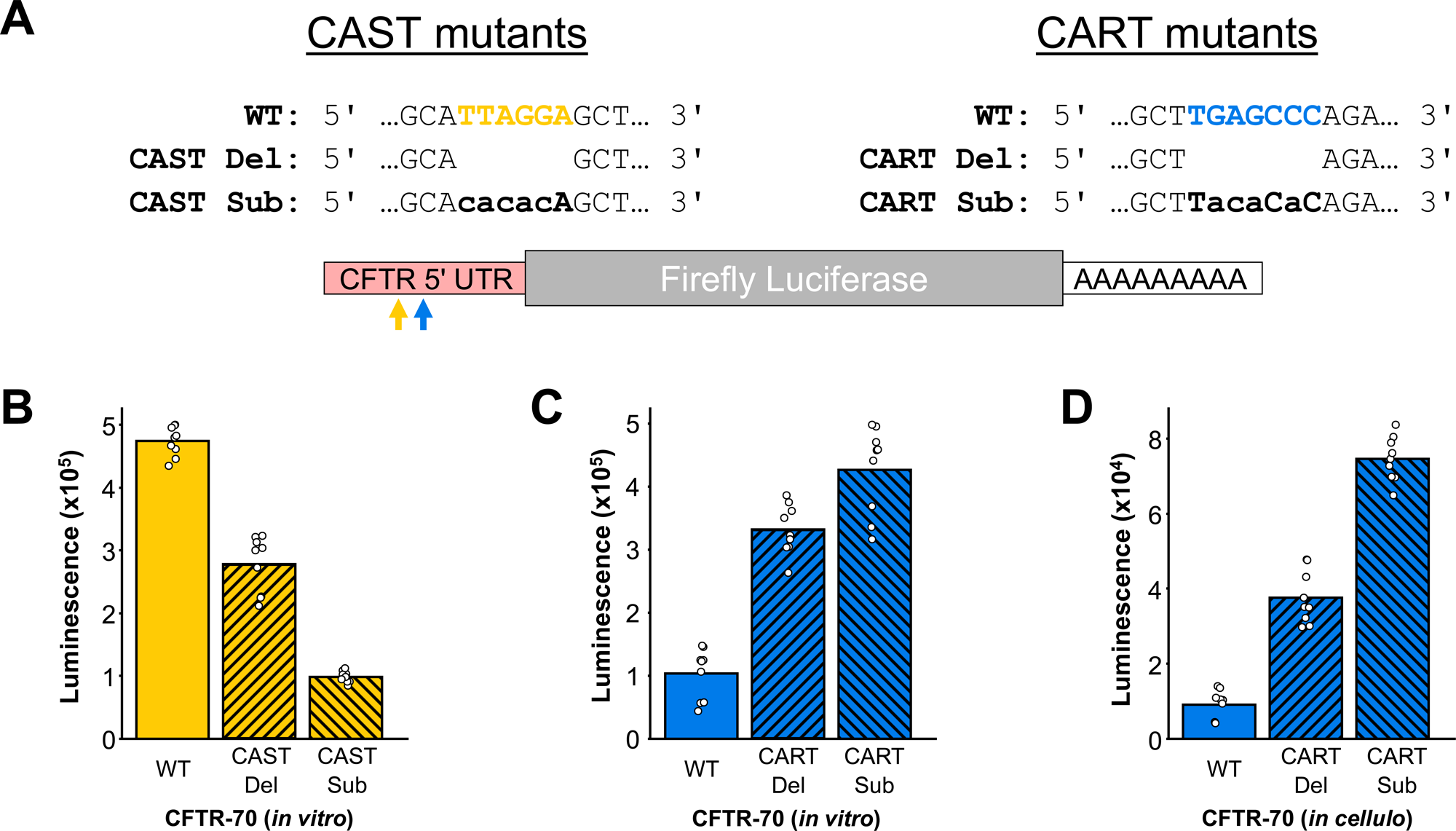
CAST and CART regulate translation of luciferase reporters. **A.** CAST and CART were disrupted in luciferase reporters containing the CFTR 5’ UTR by either deletion (Del) or substitution with CA repeats (Sub). **B-D.** *In vitro* (B, C) and *in cellulo* (D) translation assays of CAST (B) and CART (C, D) disruptions in HBE lysate (*in vitro*)/HBE cells (*in cellulo*). Bars represent averages among nine biological replicates.

### Identifying novel translational control elements in the CFTR 5’ UTR

To identify novel translational control elements, we looked for regions in the scanning deletion series that showed the highest or lowest relative RRS. As expected, the 6-nt scanning deletions had variable effects on RRS depending on the position being deleted (Fig. 4C). Generally, deletions at the 5’ and 3’ ends of the CFTR 5’ UTR tend to decrease RRS compared to the WT, which is consistent with the proposed stimulatory secondary structure (Fig. 3B). Conversely, deletions to a central region of the CFTR 5’ UTR tend to increase ribosome recruitment.

We initially focused on two regions: positions -53 to -48, where consecutive deletions decreased RRS, suggesting disruption of a potential enhancer, and positions -44 to -38 where consecutive deletions increase RRS, consistent with loss of a potential repressor (Fig. 4C). We note that while deletions near the -30-nt position increase RRS, they are not associated with an increase in RNA stability (Supp. Fig. 3A, see description of Stability Score (SS) in subsequent section). By identifying the sequence in which at least 1-nt is disrupted by all indicated deletions, we defined TTAGGA as CFTR 5’ UTR-Associated Stimulator of Translation (CAST) and TGAGCCC as CFTR 5’ UTR-Associated Repressor of Translation (CART). To investigate whether CAST and CART regulate translation, we generated luciferase reporters containing the CFTR 5’ UTR with CAST/CART either deleted or replaced by CA repeats (Fig. 5A). Both methods of removing CAST resulted in a significant decrease in protein output in an *in vitro* translation assay (Fig. 5B), consistent with CAST acting as a translational enhancer. Disruption of CART resulted in a significant increase in protein output in both *in vitro* (Fig. 5C) and *in cellulo* (Fig. 5D) translation assays, supporting that CART acts as a translational repressor. Overall, this shows our deletion strategy can be used to identify 5’ UTR regions containing translational control elements.

### CART does not regulate inherent mRNA stability

We prioritized characterization of the CART motif because as a repressor, it is most amenable to disruption to ultimately increase CFTR protein levels. One mechanism by which CART could regulate translational output is by decreasing RNA stability. We calculated a stability score (SS) by comparing the RNA abundance in the final ribosome recruitment reaction relative to the input RNA abundance. The SS reflects the relative stability of each transcript in the pool with higher values indicating higher stability. However, we note that mRNA stability is often coupled to its translational status, so we see similar effects on RRS and SS for given sequence disruptions (Supp. Fig. 3, 4). In the region containing CART in CFTR-70, we see that deletions increase SS as well as RRS (Supp. Fig. 3A, Fig. 4C). To decouple the effects on translation and stability and determine whether CART regulates the inherent stability of an mRNA, we incubated mRNAs containing the WT CFTR 5’ UTR with or without CART disruptions with ribosome-depleted HBE cell lysate and monitored RNA levels at different timepoints. We found that in the absence of translation, CART has no effect on the inherent stability of our mRNA reporters (Supp. Fig. 5), suggesting it is acting primarily at the level of mRNA translation.

### CART is enriched in membrane-protein encoding transcripts and acts broadly as a repressor

We next explored whether CART is specific to CFTR or may act more broadly. Although 5’ UTRs tend to exhibit low sequence conservation^39^, this sequence shows higher conservation than the surrounding sequence among primates, suggesting it may play an important regulatory role (Supp. Fig. 6). We investigated how prevalent CART is among other human 5’ UTRs and found that 3% of canonical human 5’ UTRs contain at least one instance of CART (Fig. 6A). Because we originally found this element in CFTR, which is a membrane protein, we investigated whether genes with CART in their 5’ UTR are enriched for membrane proteins. Indeed, CART-containing 5’ UTRs are enriched for genes encoding membrane proteins (Fig. 6B). Additionally, GO analysis of CART-containing genes revealed “regulation of monoatomic ion transmembrane transport” as the most significantly enriched term (Fig. 6C), suggesting this feature might be common to and regulate translation of other ion channels/transporters like CFTR.

**Figure 6.**
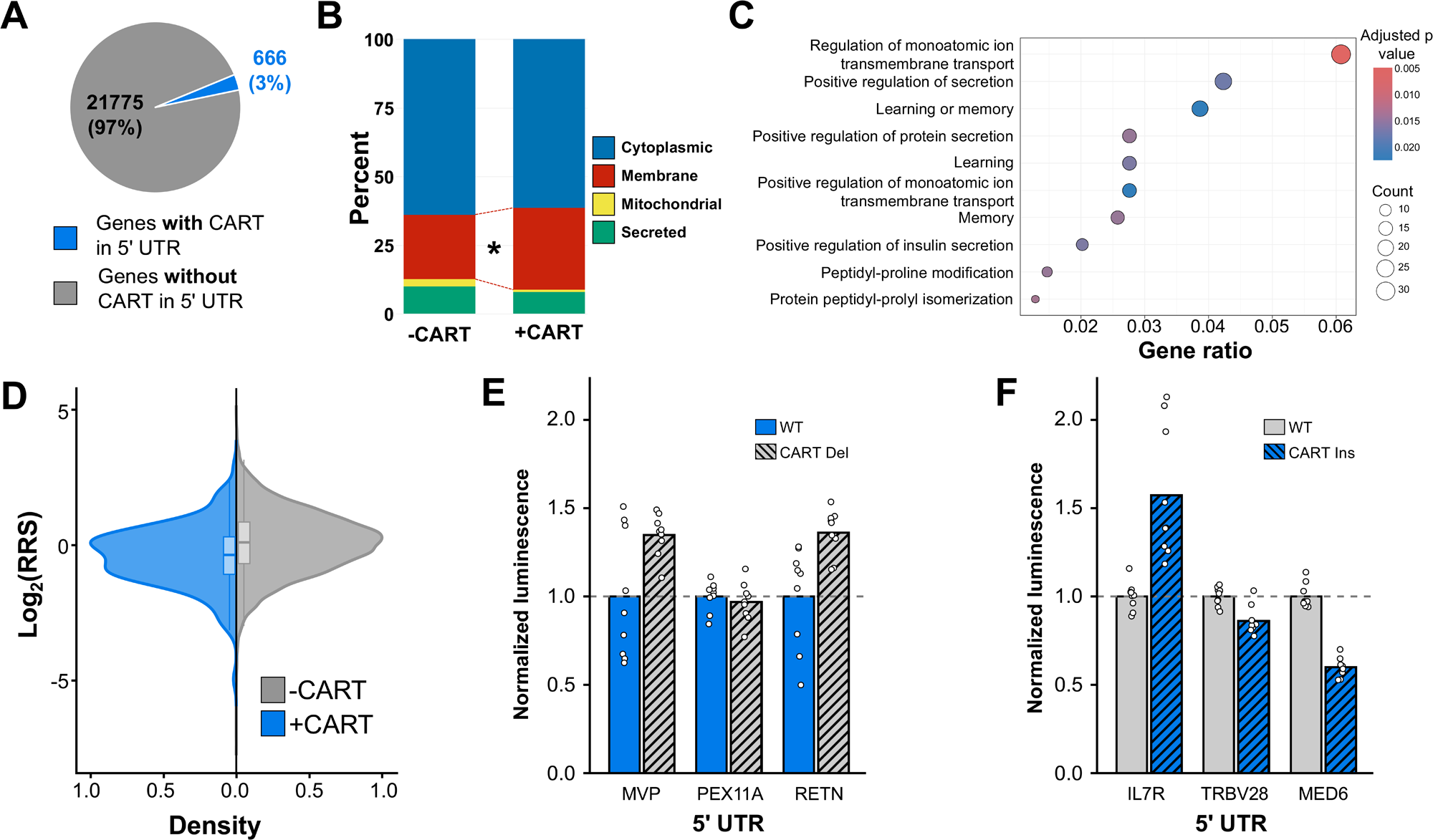
CART is found in many human 5’ UTRs and is globally associated with low ribosome recruitment and translation. **A.** Canonical human 5’ UTR sequences were obtained from Ensembl and directly searched for CART (TGAGCCC). **B.** CART-containing 5’ UTRs were subset according to the type of protein they encode (cytoplasmic, membrane, secreted, mitochondrial). P-value = 7.96E-05 (two proportion z test). **C.** GO Biological Function analysis was performed on human genes containing CART in their 5’ UTR. **D.** Distribution of RRS from a diverse pool of human 5’ UTRs was plotted for 5’ UTRs that contain or do not contain CART. P-value = 1.10E-07 (Wilcoxen rank sum test). N = 169 (+CART), N = 17981 (-CART). This assay was performed in HeLa lysate. **E.** *In vitro* translation assays were performed on luciferase reporters utilizing human 5’ UTRs containing CART and mutants which delete CART. Assays were performed using HBE lysate. Bars represent the average of nine biological replicates. **F.** *In vitro* translation assays were performed on luciferase reporters utilizing human 5’ UTRs that do not contain CART and mutants which substitute CART in the middle of the 5’ UTR. Assays were performed using HBE lysate. Bars represent the average of nine biological replicates.

To investigate whether CART acts as a translational repressor in other human 5’ UTRs, we examined DART data we collected on a pool of diverse human 5’ UTRs in HeLa lysate. We see that the presence of CART is associated with globally decreased ribosome recruitment, suggesting it functions as a translational repressor in many 5’ UTR contexts (Fig. 6D). To determine if this global reduction in RRS by the presence of CART has a similar effect on protein production, we compared protein output from luciferase reporters containing human 5’ UTRs that naturally contain CART to mutants that delete CART from the 5’ UTR. We similarly compared reporters with human 5’ UTRs that lack CART to mutants which insert CART in the middle of the 5’ UTR. We found that deletion of the CART motif increased translation for two of the three tested constructs (Fig. 6E), and insertion of CART decreased translation for two of the three tested constructs (Fig. 6F). Together, these data support that CART acts broadly as a translational repressor, although the magnitude of its impact is context dependent.

Overall, we outline a strategy to systematically identify 5’ UTR features that regulate translation of a gene of interest. Using the CFTR 5’ UTR, we demonstrated that this strategy detects known 5’ UTR regulatory elements like uAUGs and structures. We also showed using a large set of targeted mutations that we can identify regions of a given 5’ UTR harboring repressive and stimulatory features that cannot be predicted from sequence alone. We used these mutants to identify a translational stimulator (CAST) and repressor (CART) in the CFTR 5’ UTR, with CART being an attractive therapeutic target to increase CFTR protein production. Finally, we showed CART is present in many other human 5’ UTRs from related genes and may more broadly regulate translation.

## Discussion

Identification of targetable translational control elements has potential therapeutic benefit for many diseases. Here, we demonstrate a strategy to identify targetable translational control elements in a 5’ UTR using CFTR as an example, as modulating translation of this protein is expected to be therapeutically valuable^31–34^. We measured ribosome recruitment to a pool of systematically mutated CFTR 5’ UTRs and successfully identified features regulating translation. This strategy enabled us to test the effects of predicted regulatory features, like uAUGs and secondary structure, as well as identify regions containing previously unknown regulatory features that cannot be predicted from sequence alone. This strategy ultimately identified differences in translation of tissue-specific CFTR 5’ UTR isoforms, regulatory secondary structures, and previously unknown sequence-based enhancers and repressors that could be targeted therapeutically to modulate CFTR translation.

DART revealed differences in ribosome recruitment between tissue-specific CFTR 5’ UTR isoforms. We found that CFTR-132 exhibits a relatively high RRS due to the presence of a functional uAUG, even though the uAUG is out-of-frame and represses protein output from the main AUG in a translation assay. Although this could lead to ambiguity when comparing RRS between transcripts with different amounts of uAUGs, we were intrigued by the possibility of identifying functional uAUGs using DART. We analyzed ribosome recruitment data obtained on a pool of diverse human 5’ UTRs that includes a pathogenic 5’ UTR mutant of POMC (-11C>A), which forms an out-of-frame uAUG and is known to repress translation at the main AUG^40^. Presence of the functional uAUG in the pathogenic POMC mutant increased ribosome recruitment but decreased protein output by an *in vitro* translation assay (Supp. Fig. 7), supporting that DART can be used to identify functional uAUGs in multiple 5’ UTR contexts.

We note that translational output of CFTR-132 was previously reported to be about twice that of CFTR-70 using a plasmid-based translation assay in HT29 and HEK293 cells^36^, which is inconsistent with our findings that CFTR-70 generates about twice as much protein as CFTR-132. In our assay, we prepared capped mRNA *in vitro* and mixed it with translationally active cytoplasmic HBE lysate. We use this strategy to avoid ambiguity: changes to the 5’ UTR are inherently close to the promotor region and can therefore influence expression, transcription start site selection, and splicing. By using mRNA reporters in an *in vitro* system, we eliminate many of these confounding variables. However, this strategy comes with its own caveats: the removal of cell components like membranes during lysate generation and lack of RNA modifications and RBPs on the *in vitro* transcribed mRNA may affect translational regulation.

To investigate the role of 5’ UTR secondary structures on translation of CFTR, we measured ribosome recruitment to a mutational series targeting predicted secondary structures in the CFTR 5’ UTR. In CFTR-70, we found that the most highly predicted secondary structure (stem loop 1) enhances ribosome recruitment. This finding is intriguing as secondary structures in 5’ UTRs typically repress translation initiation^16–18^, and it has been previously suggested that this stem actually represses translation^36^. Mechanistically, this structure could be enhancing translation through direct recruitment of ribosomes (IRES) or initiation factors (CITE), although IRES are typically larger and we see low translational output of uncapped CFTR-70 luciferase reporters (Supp. Fig. 8). It is also possible that the structure favors recruitment of helicases or other RNA binding proteins that promote translation overall. Further experiments are required to fully elucidate the mechanism at work.

We next systematically identified targetable regulatory features in the CFTR 5’ UTR by measuring ribosome recruitment to a large set of targeted mutations across the entire length of the CFTR-70 5’ UTR. We expect this approach to be broadly applicable to other genes where manipulating translational output may be therapeutically beneficial. Although the scanning substitution strategy made identification of regulatory elements difficult, it highlights the large effect a small sequence in a 5’ UTR can have on ribosome recruitment and translation. We hypothesize these data could be used to identify robust translational enhancers and repressors that are likely present in human 5’ UTRs but remain completely uncharacterized.

Overall, we found that the 6-nt scanning reported more directly on the endogenous sequence elements, which we used for preliminary identification of stimulatory and repressive elements CAST and CART. We used translation assays to show that CAST enhances and CART represses translational output. Interestingly, CART is found in many human 5’ UTRs and is enriched for membrane proteins related to CFTR: monoatomic ion transporters. A thorough investigation is warranted into the relationship between these genes and CFTR and whether CART is co-regulating ion transporters as a group. Our studies show the magnitude of CART’s repressive effect is somewhat context dependent, therefore direct testing of the motif is required.

Finally, our data suggest CART represents a promising mutant-agnostic therapeutic target to enhance translation of CFTR. The exact mechanism of repression will dictate what therapeutic options are most likely to be successful. However, due to the prevalence and portability of the CART motif we hypothesize that it is likely acting through a protein binding partner rather than a specific secondary structure. If so, it represents an attractive target for steric blocking by antisense oligonucleotides. Looking forward, we believe our pipeline will be broadly useful for identifying new 5’ UTR regulatory elements that can be used to manipulate translation of other disease-relevant genes.

## Materials and Methods

### Direct Analysis of Ribosome Targeting

#### Experimental setup

BP1 and RN1 ssDNA oligo pools were ordered from Twist Bioscience and used to prepare pooled mRNA as previously described^25,41^ with the following modifications: Oligos contain (from 5’ to 3’) the T7 promoter (CTAATACGACTCACTATA<u>G</u>GG), a variable 5’ UTR sequence with variable length of downstream coding sequence, a randomized 10-nt barcode, and a reverse transcription primer binding site (TGGAATTCTCGGGTGCCAAG). In BP1, five copies of each of the three WT CFTR 5’ UTR isoforms (CFTR-70, CFTR-107, CFTR-132) were included, which are identical except for the 10-nt barcode.

Oligo pools were PCR amplified as described previously^25,41^. RN1 was purified using the Zymo DNA Clean & Concentrator-5 (Zymo Research, D4004) according to the manufacturer’s instructions, and the BP1 pool was purified by phenol:chloroform extraction and isopropanol precipitation. Amplified pools were *in vitro* transcribed by assembling the following reaction: 0.8115 pmol (for the RN1 pool) or 10.82 pmol (for the BP1 pool), 40 mM HEPES-KOH pH = 7.5, 10 mM NaOAc, 75 mM Mg(OAc)_2_, 0.2 mM spermidine, 40 mM DTT, 10 mM ATP, 10 mM CTP, 10 mM GTP, 10 mM UTP, 2000 U/mL RNasin Plus Ribonuclease Inhibitor (Promega Corporation, N2611), and homemade T7 RNA polymerase mutant^42^. *In vitro* transcriptions were incubated at 37°C overnight in a thermal cycler. Transcribed RNA was gel purified as described^25,41^.

A 7-methyl guanosine cap was added to the transcripts in the BP1 pool using the Faustovirus capping system (New England Biolabs, M2081) according to the manufacturer’s instructions. Transcripts in the RN1 pool were capped using the Vaccinia capping system (New England Biolabs, M2080) according to the manufacturer’s instructions. The capped BP1 pool was gel purified on an 8% polyacrylamide gel as described^25,41^, and the RN1 pool was purified by phenol:chloroform extraction and isopropanol precipitation.

#### Ribosome recruitment reaction

Ribosome recruitment reactions were performed in triplicate by assembling the following reaction: 6.24 pmol (for the BP1 pool) or 15.60 pmol (for the RN1 pool), 16 mM HEPES-KOH pH = 7.6, 50 mM KOAc, 2.5 mM Mg(OAc)_2_, 100 μM spermidine, 20 mM creatine phosphate, 100 μg/mL creatine phosphokinase (Sigma-Aldrich, C3755-500UN), 400 U/mL RNasin Plus Ribonuclease Inhibitor (Promega Corporation, N2611), and 100 μg/mL cycloheximide in 86.7% HBE lysate (for the BP1 pool, see “Generating lysates”) or commercial HeLa lysate (for the RN1 pool (Ipracel, CC-01-40-50)). Commercial HeLa cell lysate was clarified by centrifugation at 12,000 x g for 2 minutes at 4°C. Reactions were prepared on ice and then incubated at 37°C for 30 minutes in a thermal cycler. Following incubation, 13% of each reaction was removed and flash frozen for subsequent RNA extraction/library preparation (this is the incubated RNA sample used for calculating stability scores). The remainder of the reactions were loaded onto 10-50% linear sucrose gradients (20 mM HEPES-KOH pH=7.6, 100 mM KCl, 5 mM MgCl_2_, 1 mM DTT, 100 μg/mL cycloheximide) and centrifuged at 35000 rpm for 3 hours at 4°C in an SW41 rotor (Beckman Coulter). Following centrifugation, gradients were fractionated using a piston gradient fractionator (BioComp). Fractions corresponding to the 80s peak were combined and total RNA was extracted by phenol:chloroform extraction and isopropanol precipitation.

Library preparation was performed on 80s-bound RNA, incubated RNA, and the transcribed RNA pool (called RNA-only input) as previously described^25,41^, except no biotin pulldown was performed (pools were not biotinylated). For reverse transcription of the RNA-only input, 0.156 pmol (for the BP1 pool) or 15.60 pmol (for the RN1 pool) was used.

#### Data processing

Libraries underwent paired-end sequencing on an Illumina NovaSeqXPlus (for the BP1 pool) or Illumina NextSeq 2000 (for the RN1 pool). Sequencing and demultiplexing were performed by the Advanced Genomic Core at the University of Michigan. Raw sequencing data was assessed using FastQC. Adapters and low-quality regions of reads were trimmed using fastp and UMIs (10-nt located on read 2) were extracted and annotated. Trimmed reads were aligned to a custom reference containing all transcripts in the pool using STAR aligner. Per-reference counts were generated from BAM files using pileup.sh from BBMap. The counts for each transcript were normalized by transcripts per million (TPM) for each library. Ribosome recruitment scores (RRS) were calculated by dividing the normalized read count in the 80s library by the normalized read count in the RNA-only input library. Stability scores (SS) were calculated by dividing the count in the incubated RNA library by the count in the RNA-only input library.

Six-nucleotide scanning deletions and substitutions were plotted as the log_2_ of the fold change in the RRS of the mutant (either a deletion or substitution) to the average of the five copies of a given WT transcript (for each WT CFTR 5’ UTR, 5 individually barcoded transcripts were included in the pool). Relative RRS values from scanning deletions which form an AUG were replaced with transcripts which incorporate the 6-nt deletion and further mutate the AUG to AGT to avoid ambiguous scores from multiple start sites. Scanning substitutions were designed such that they never form an AUG. All RRS data was plotted in R^43^.

### Generating lysates

Cell lysates used for DART and *in vitro* translation assays were generated from human bronchial epithelial cells (16HBE14o-, indicated here as HBE). HBE cells were grown in DMEM (Gibco™, 11965126) supplemented with 10% FBS (Biowest, S1620) and 1 x Penicillin-Streptomycin Solution (Corning, 30-002-CI) on collagen coated plates. To make the coating solution, a mixture of 0.01% Bovine Serum Albumin (Gibco™, 15260-037), 1% type 1 bovine collagen solution (Advanced BioMatrix, 5005), and 10 μg/mL of human fibronectin (Gibco™, 33016-015) was prepared in LHC-8 basal medium (Gibco™, 12677-027). To each 15 cm plate, 5 mL of coating solution was added and allowed to incubate at 37°C for 2-3 hours. After the incubation, the coating solution was removed, and plates were stored at 4C. Plates were used within 1 month of coating.

Batches of 10-20 15 cm TC-treated plates of cells were grown to approximately 90% confluence, washed with 1 x PBS (Corning, 21-040-CV), and harvested by trypsinization (Corning, 25-053-CI). Cells were counted on a Countess 3 Automated Cell Counter (Thermo Fisher Scientific) and resuspended in 1 mL of hypotonic buffer (10 mM HEPES-KOH pH = 7.6, 10 mM KOAc, 0.5 mM Mg(OAc)_2_, 5 mM DTT, 1x protease inhibitor cocktail (Roche, 11836170001)) for every 100-200 million cells. Cells were then lysed by centrifugation in a ZentriMix 380 R dual centrifuge (Hettich) in 2 mL tubes filled with 500 μL cells and under the following conditions: 1000 rpm, 4 minutes, -5°C. Cells can alternatively be lysed using other standard mechanical lysis methods like Dounce homogenization, passage through a thin needle, or sonication. Cells were stained with Trypan Blue (Invitrogen™, T10282) after dual centrifugation and visualized by confocal microscopy to ensure greater than 50% lysis. Lysates were clarified by centrifugation at 13000 x g for 10 minutes at 4°C in a microcentrifuge. Supernatant was aliquoted and flash frozen in liquid nitrogen.

### Luciferase reporter design and preparation of mRNA

Plasmids encoding firefly luciferase downstream of 5’ UTR sequences of interest were made either by isothermal assembly or site-directed mutagenesis with partially overlapping primers. These plasmids contain the T7 promotor sequence followed by the 5’ UTR of interest, firefly luciferase coding sequence, a short synthetic 3’ UTR, and a 50-nt poly-A sequence. These components are within a standard high copy number cloning vector. Below are relevant sequences used for reporter design:

5’ UTR sequences:

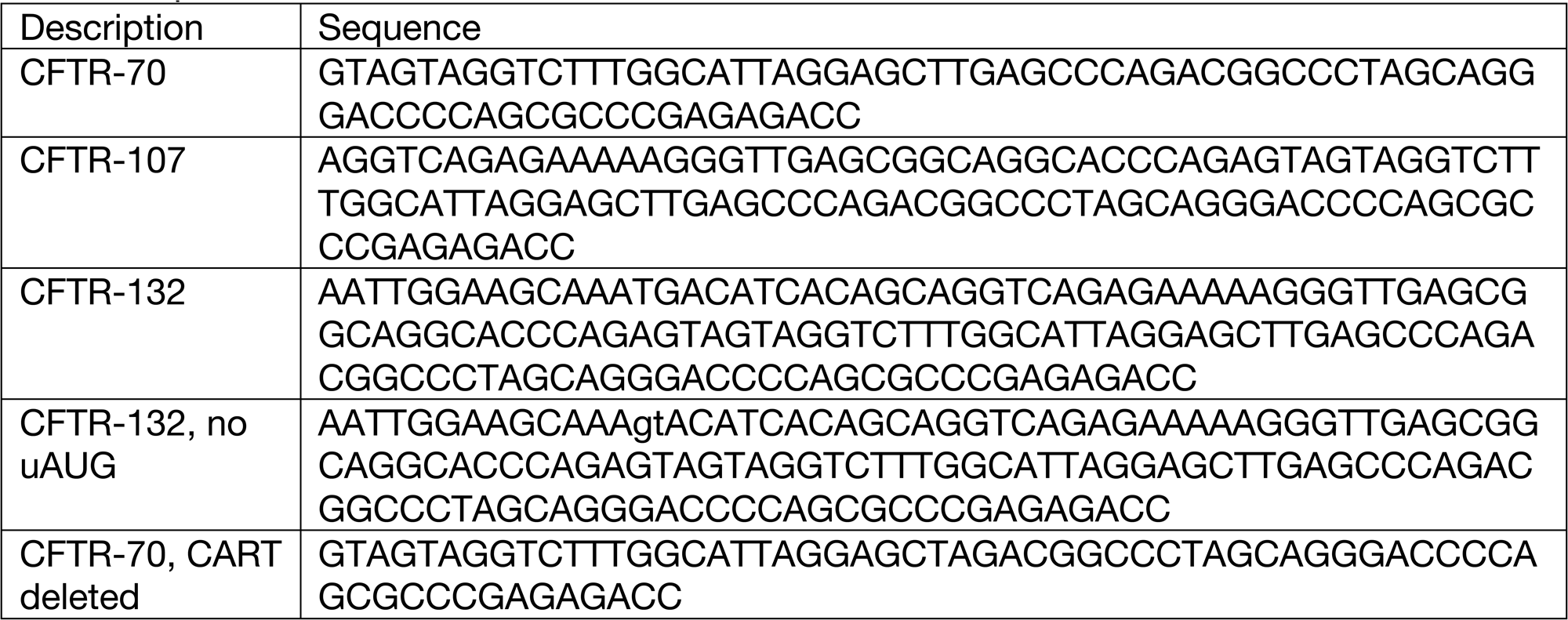
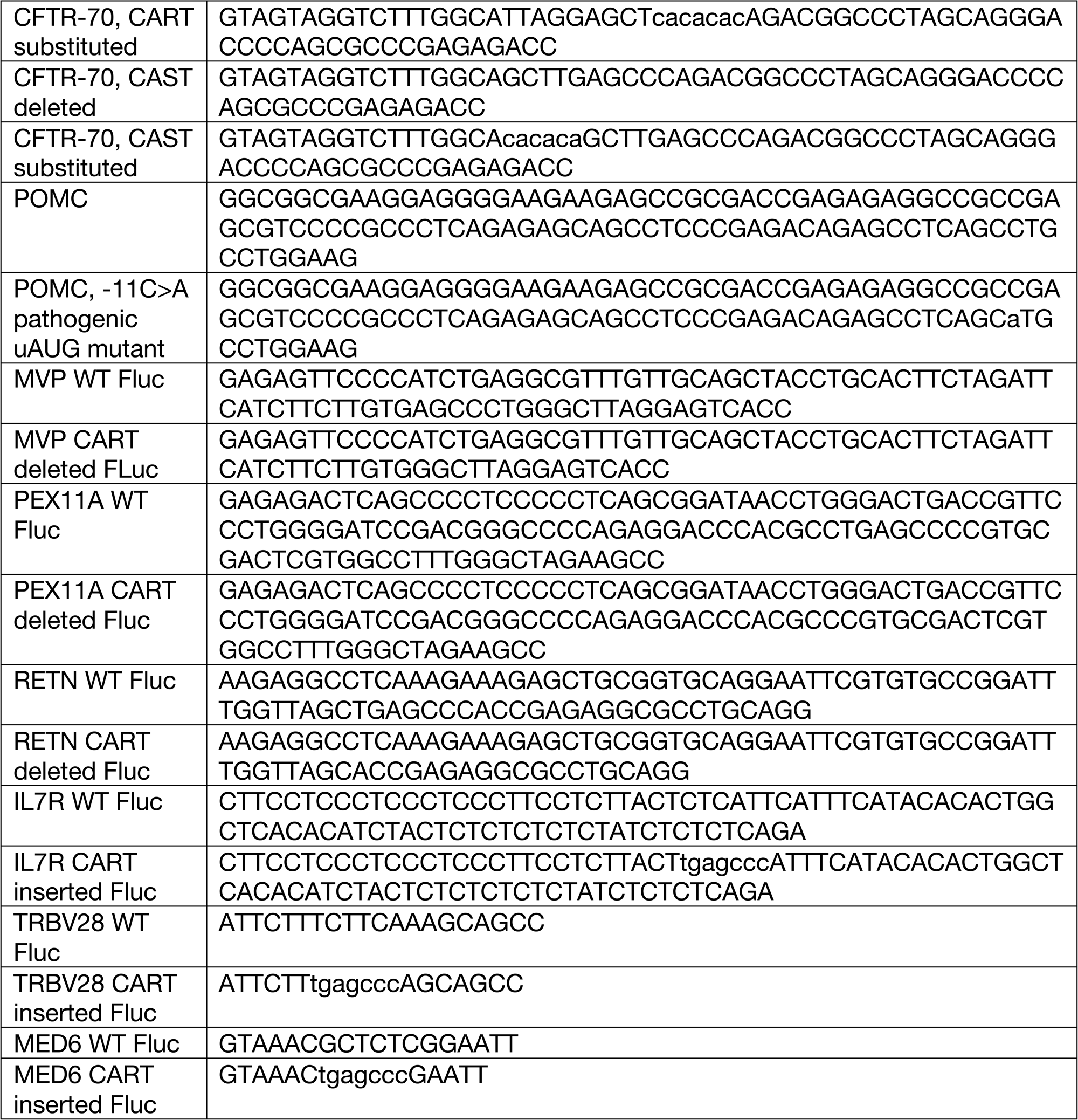

T7 promotor sequence: TAATACGACTCACTATA<u>G</u>GG

Minimal 3’ UTR: TAGATCCACTAGTTCTAGAGCGGCCGCCACCGCGGTGGAGGGGTAC

### Preparation of luciferase reporter mRNA

Plasmids encoding firefly luciferase (see Luciferase reporter design and preparation of mRNA) were linearized using a single Ecl136II site according to the manufacturer’s instructions (Thermo Scientific™, FERFD0254). Linearized DNA was purified using the Zymo DNA Clean & Concentrator-5 kit (Zymo Research, D4004) according to the manufacturer’s instructions. mRNA was *in vitro* transcribed by diluting 0.1642-1.642 pmol of linearized DNA in the following components: 40 mM HEPES-KOH pH = 7.5, 10 mM NaOAc, 75 mM Mg(OAc)_2_, 0.2 mM spermidine, 40 mM DTT, 10 mM ATP, 10 mM CTP, 10 mM GTP, 10 mM UTP, 2000 U/mL RNasin Plus Ribonuclease Inhibitor (Promega Corporation, N2611), and homemade T7 RNA polymerase mutant^42^. *In vitro* transcriptions were incubated at 37°C for 3 hours in a thermal cycler. Transcribed RNA was purified using the Zymo RNA Clean and Concentrator-25 kit (Zymo Research, R1017) according to the manufacturer’s instructions. Purified RNA was *in vitro* capped using the FCE capping system (New England Biolabs, M2081) according to manufacturer’s instructions. Capped RNAs were purified using the Zymo RNA Clean and Concentrator-25 kit (Zymo Research, R1017) according to the manufacturer’s instructions and quantified using the Qubit RNA Broad-Range Assay Kit (Invitrogen™, Q10210). RNAs were diluted to 82-164 nM stocks for translation assays.

Equal volumes of diluted mRNAs were run on a glyoxal gel to ensure quality of mRNA reporters before use in *in vitro*/*cellulo* translation assays. Band intensity for each RNA stock was also quantified from the gel and used to input-normalize luminescence values from translation assays.

### *In vitro* translation assays

*In vitro* translation assays were performed by preparing the following reaction: 16.4 nM mRNA reporter (see “Luciferase reporter design and preparation of mRNA”), 16 mM HEPES-KOH pH = 7.6, 50 mM KOAc, 2.5 mM Mg(OAc)_2_, 100 μM spermidine, 20 mM creatine phosphate, and 100 μg/mL creatine phosphokinase (Sigma-Aldrich, C3755-500UN), and 400 U/mL RNasin Plus Ribonuclease Inhibitor (Promega Corporation, N2611) in 50% HBE lysate (see “Generating lysates”) or 50% commercial HeLa lysate (Ipracell, CC-01-40-50). Reactions were prepared on ice and then incubated at 37°C for 30 minutes in a thermal cycler. After the incubation, reactions were moved to ice and diluted two-fold by adding an equal volume of 2x Passive Lysis Buffer (Promega Corporation, E1941). 3/4 volumes of this mixture were added to wells of a white round bottom 96-well plate (Corning™, 07-200-336) and diluted two-fold by mixing with an equal volume of BrightGlo reagent (Promega Corporation, E2610) which was previously added to the plate. After incubating at room temperature for 3 minutes, the plate was centrifuged in a tabletop centrifuge (3000 x g for 3 minutes) to remove bubbles and luminescence was measured on a Spark plate reader (Tecan) using the following settings: mode: luminescence, attenuation: none, settle time: 0, integration time (ms): 1000, output: counts/s, temperature: 21°C. Data were normalized to RNA input as described in “Preparation of luciferase reporter mRNA”. Data were plotted in R^43^.

### *In cellulo* translation assays

*In cellulo* translation assays were performed by transfecting 16HBE14o-cells with .164 pmol of luciferase reporter (see “Luciferase reporter design and preparation of mRNA”). HBE cells were grown in DMEM (supplemented with 10% FBS (Biowest, S1620) and 1x Penicillin-Streptomycin Solution (Corning, 30-002-CI)) and seeded at 30,000 cells per well of a clear bottom tissue culture-treated 96-well plate (Corning™, 07-200-565) 24 hours before transfection. HBE cells were transfected with the indicated quantity of luciferase reporter using Lipofectamine MessengerMAX Transfection Reagent (Invitrogen™, LMRNA003) according to the manufacturer’s instructions. At approximately 24 hours post transfection, media was removed and 40 μL of 1x Cell Culture Lysis Reagent (Promega Corporation, E1531) was added to each well. Plates were incubated for 15 minutes at room temperature with rocking. An equal volume of BrightGlo (Promega Corporation, E2610) was added directly to each well and mixed. After incubating at room temperature for 3 minutes, the plate was centrifuged in a tabletop centrifuge (3000 x g for 30 seconds) to remove bubbles and luminescence was measured on a Spark plate reader (Tecan) as described above (see “ *In vitro* translation assays”). Data were normalized to RNA input as described in “Preparation of luciferase reporter mRNA”. Data were plotted in R^43^.

### mRNA stability assay

The inherent stability of mRNAs was tested by mixing them in ribosome-depleted cell lysate, purifying total RNA, and directly visualizing RNA degradation by RNA capillary gel electrophoresis. Ribosome-depleted cell lysate was generated by first preparing HBE lysate as described above (see “Generating lysates”). This cell lysate was placed in thick wall polycarbonate tubes (Beckman Coulter, 343778) and centrifuged in a TLA120.2 rotor (Beckman Coulter) at 80000 rpm (278000 x g) for 30 minutes at 4°C (adapted from a previous protocol to remove ribosomes from rabbit reticulocyte lysate^44^). The supernatant was moved to a new polycarbonate tube and centrifuged again using the same conditions. These steps were repeated until very little/no pellet was observed (this usually takes about 3 x 30 minute centrifugations). Following centrifugation, ribosome depleted lysate was aliquoted and flash frozen in liquid nitrogen. Depletion of ribosomes was confirmed by *in vitro* translation assay using ribosome depleted lysate and the same lysate supplemented with resuspended ribosomes (data not shown).

RNA stability assays were performed by mixing mRNA reporters (see “Luciferase reporter design and preparation of mRNA”) and ribosome-depleted HBE lysate to these concentrations: 164.22 nM reporter and 80% ribosome-depleted HBE lysate. Aliquots were removed from the tubes after 0, 0.5, 1.5, and 5 minutes, added to 1.5mL tubes containing TriZol (Invitrogen™, 15596026), and immediately flash frozen in liquid nitrogen. After completion of the time course, total RNA was purified from samples according to manufacturer’s instructions.

After RNA purification, the concentration of each sample was measured using the Qubit RNA High-Sensitivity Assay Kit (Invitrogen™, Q32852) and each sample was diluted to 25 ng/μL greater than the concentration of a no RNA control. This was done so that the exogenous mRNA falls within the analytical range of the RNA capillary gel electrophoresis system. The ribosome depleted cell lysate contains small RNAs (tRNAs, 5s rRNA) which contribute substantially to the concentration of RNA in the purified samples (but do not interfere with visualization). After dilution, RNA degradation was visualized using a High Sensitivity RNA ScreenTape (Agilent Technologies, 5067-5579, 5067-5580, 5067-5581) according to the manufacturer’s instructions.

### Analysis of human 5’ UTRs

Canonical human 5’ UTR sequences were retrieved from Ensembl using the Biomart package^45,46^ in R^43^. To quantify the fraction of human genes that contain CART in the 5’ UTR, the CART sequence (TGAGCCC) was directly searched in all canonical human 5’ UTR sequences. Gene IDs from this list were queried in UniProt to obtain attributes used to further subset human genes into the class of proteins they encode. Specifically, genes containing a transit peptide were labeled “mitochondrial”, genes containing a transmembrane domain were labeled “membrane”, genes containing a signal peptide but no transmembrane domain were labeled “secreted”, and genes lacking a transit peptide, signal peptide, and transmembrane domain were labeled “cytoplasmic”. For GO Biological Process analysis, gene IDs for genes containing CART in the 5’ UTR were fed into the ClusterProfiler R package^47,48^ using the org.Hs.eg.db reference database^49^.

## ABBREVIATIONS

CF: cystic fibrosis;
CFTR: Cystic Fibrosis Transmembrane Conductance Regulator;
5’ UTR: 5’ untranslated region.

---

**Supplementary Figure 1.**
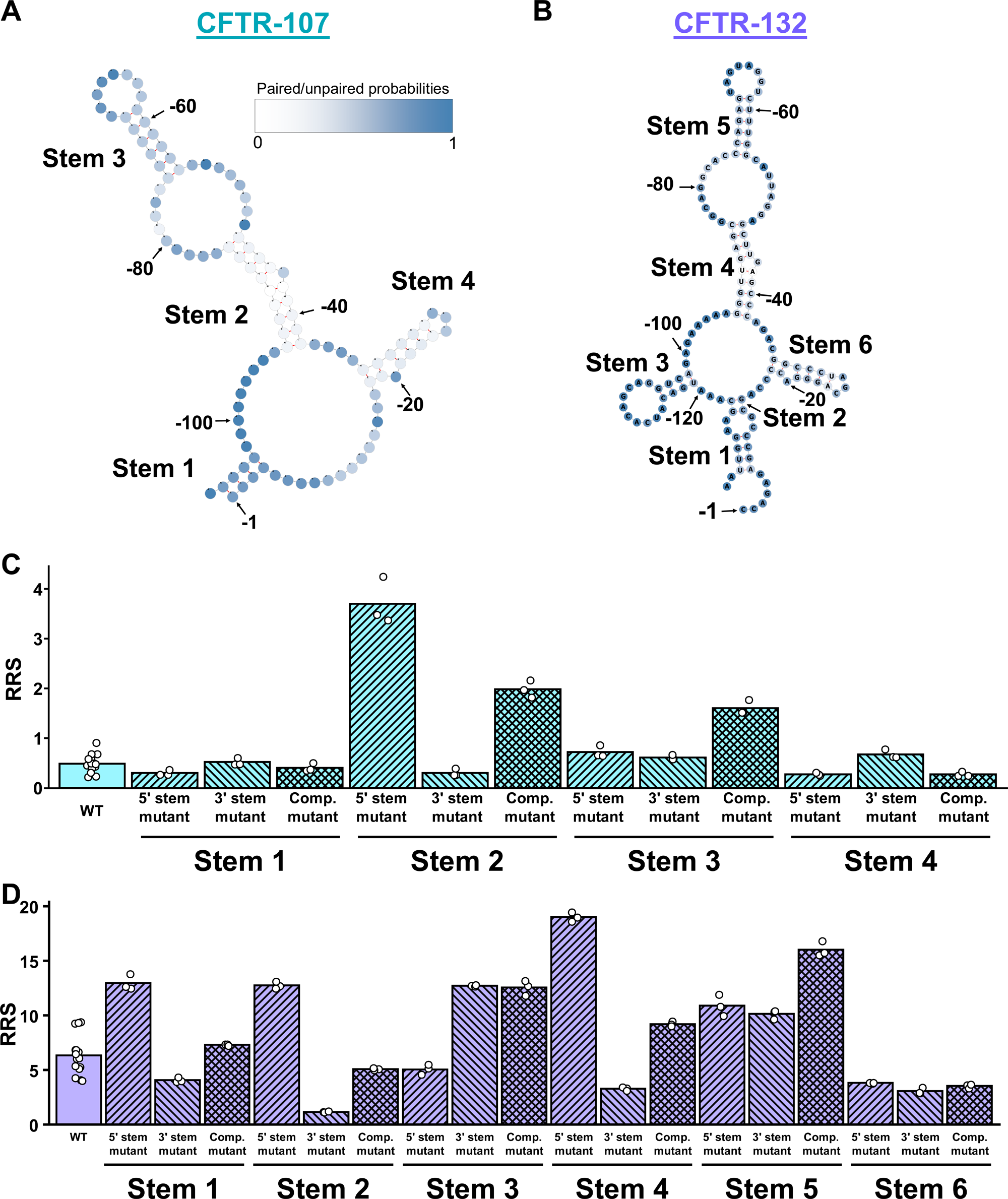
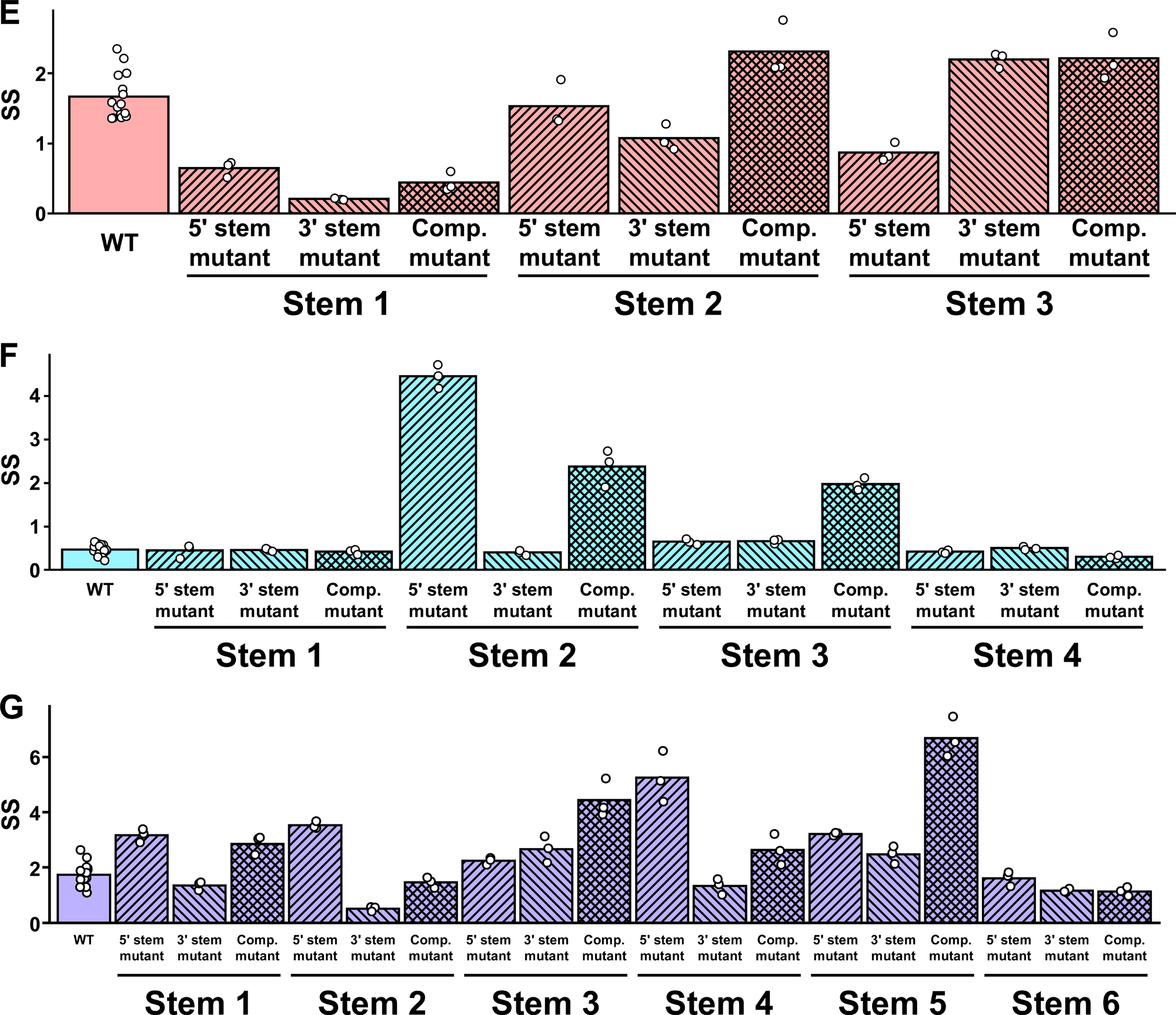
Predicted minimum free energy structures of CFTR-107 and CFTR-132 and associated DART data on structure disrupting mutants. **A, B**. Minimum free energy RNA secondary structures were predicted using the RNAfold web server for the CFTR-107 (**A**) and CFTR-132 (**B)** 5’ UTR isoforms and used to make targeted mutations to disrupt each predicted stem loop (see Fig. 3A, supplementary table 1). Nucleotides are labeled according to their position with respect to the first nucleotide of the start codon. Secondary structures were visualized using forna. **C, D.** RRS were plotted for each mutant and the WT for the CFTR-107 **(C)** and CFTR-132 **(D)** 5’ UTR isoforms. **E-G.** Stability scores (SS) were plotted for each structure disrupting mutation and the WT for the CFTR-70 **(E)**, CFTR-107 **(F)**, and CFTR-132 **(G)** isoforms. For plots C-G, bars are the average of three biological replicates (dots), except the WT which is an average of three biological replicates for five individually barcoded transcripts in the pool.

**Supplementary Figure 2.**
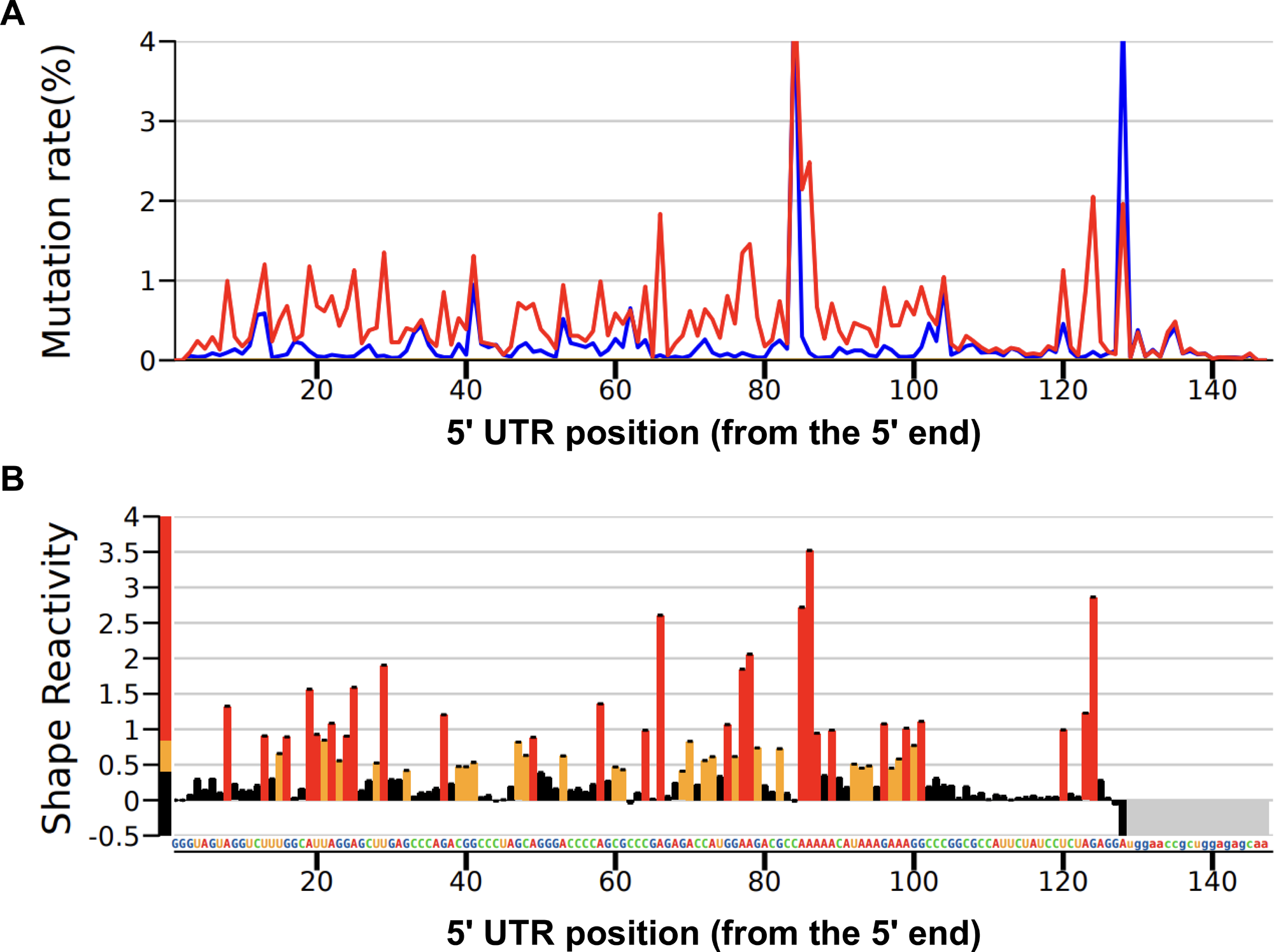
SHAPE-MaP was performed on the CFTR-70 5’ UTR firefly luciferase reporter (illustrated in figure 5A). **A.** Mutation rate was calculated for each nucleotide between positions -70 to +56 in the CFTR-70 5’ UTR firefly luciferase reporter for an untreated control (blue line) and 1M7-treated sample (red line). **B.** Mutations rates for the treated/untreated samples were used to calculate the shape reactivity. Nucleotides with shape reactivity of 0-x were labeled black (unreactive), nucleotides with shape reactivity of x-y were labeled orange (moderate reactivity), and nucleotides with shape reactivity of y and above were labeled red (high reactivity).

**Supplementary Figure 3.**
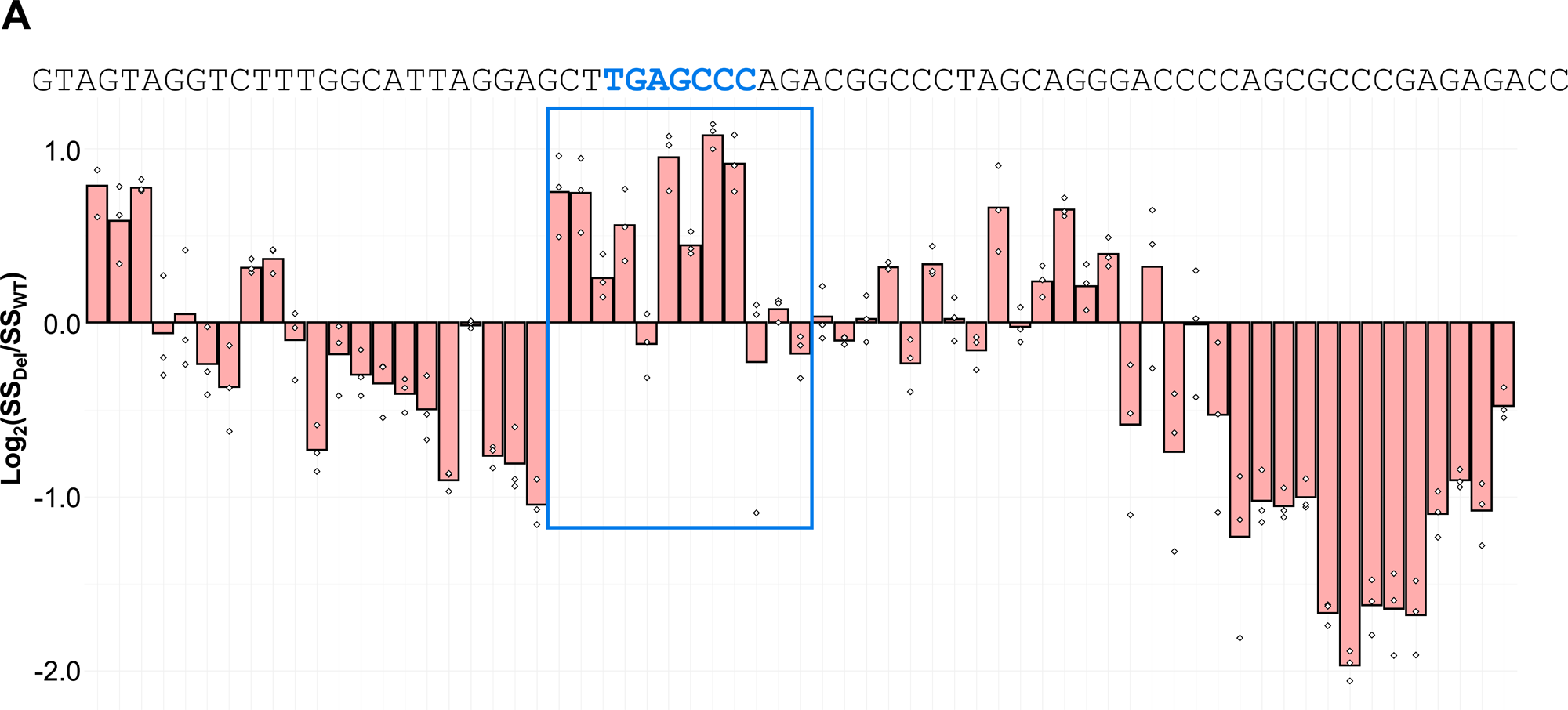
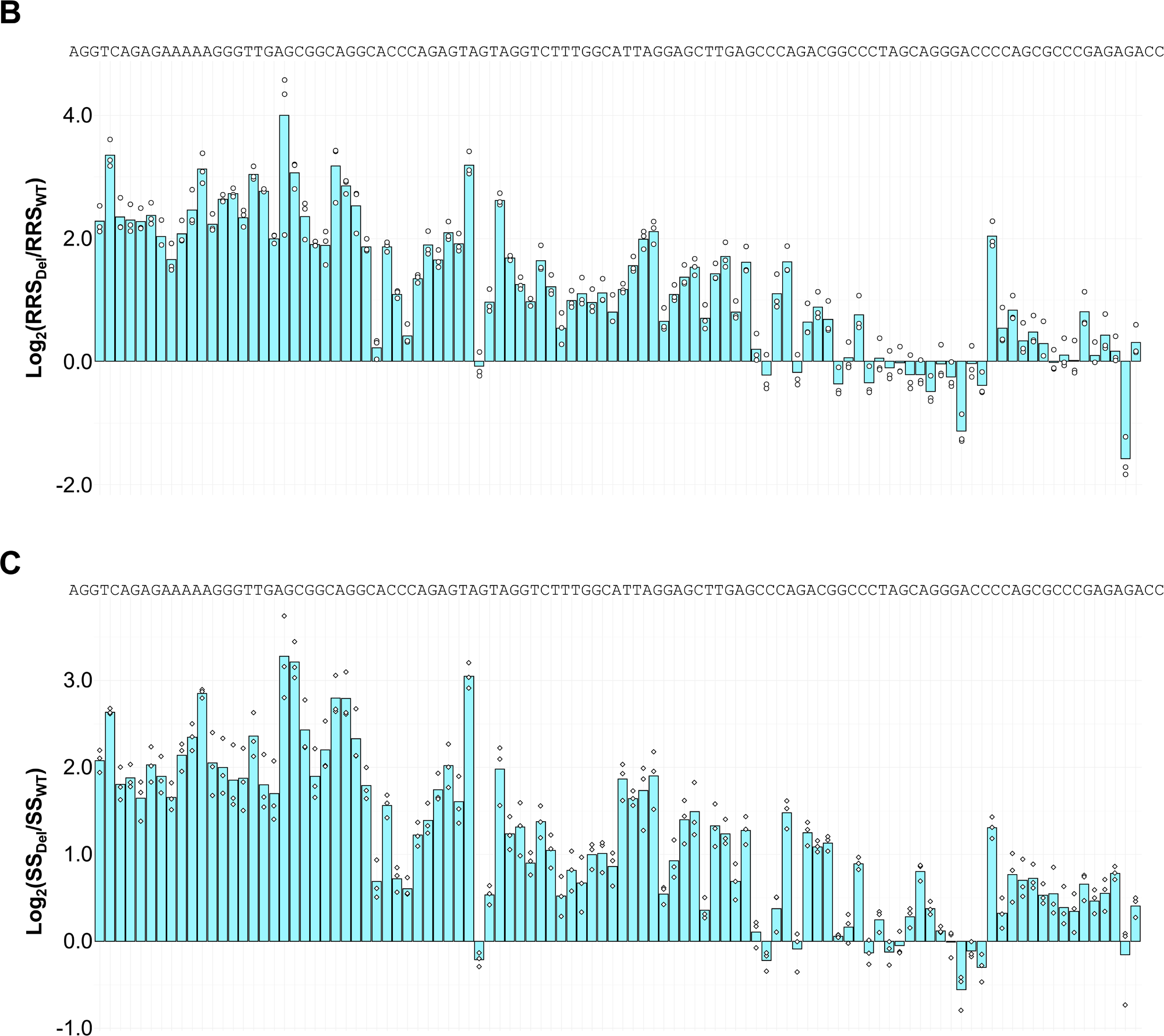
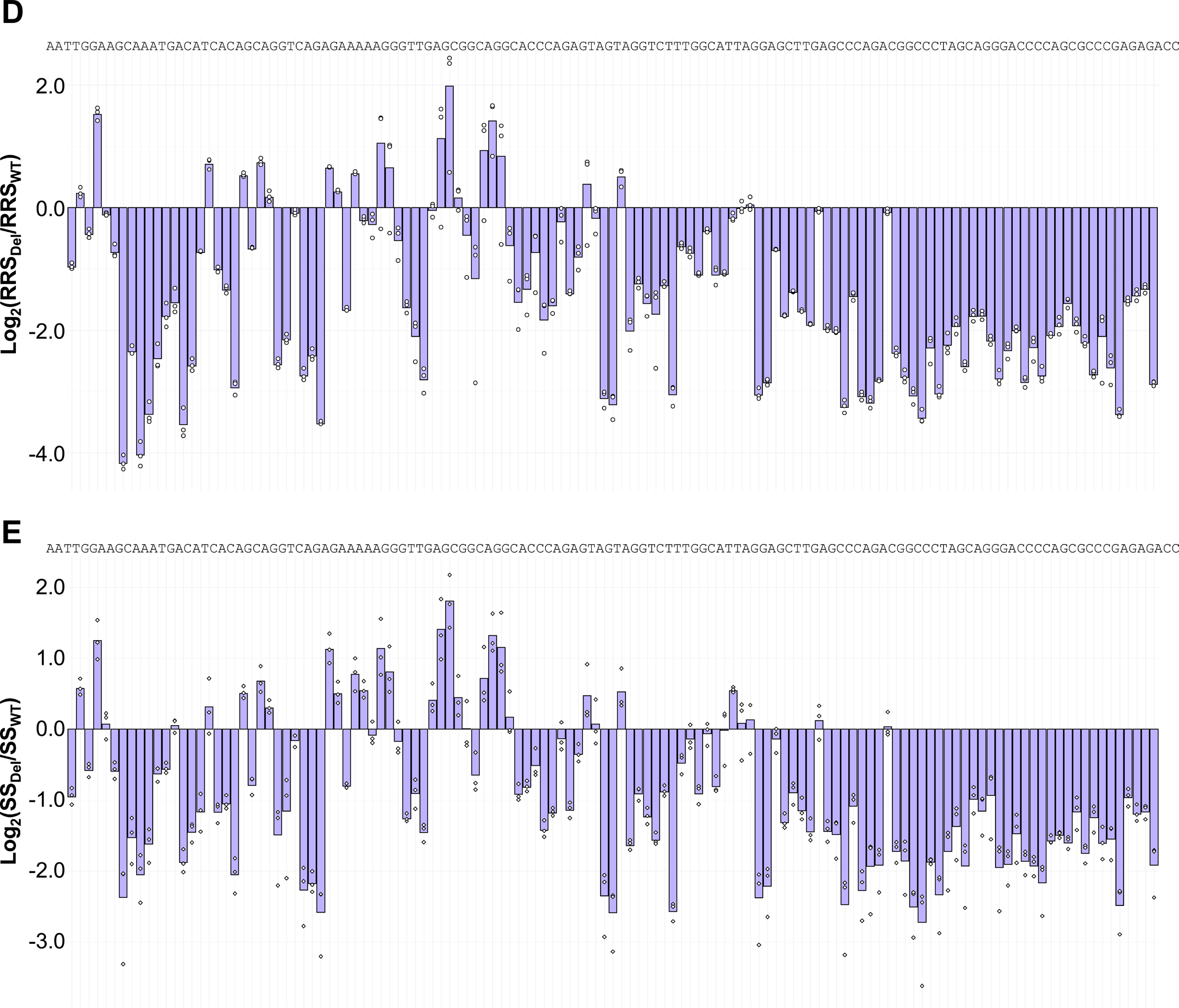
Ribosome recruitment scores (RRS) and stability scores (SS) for CFTR-70 (**A**), CFTR-107 (**B**, **C**), and CFTR-132 (**D**, **E**) scanning deletions. The log_2_ fold change (FC) of the RRS/SS relative to the WT was calculated for each 6-nt deletion. Each bar represents the average log_2_FC for a given deletion for three biological replicates (dots). Bars are centered on the 6-nt that are deleted (the WT CFTR 5’ UTR sequence is aligned on top).

**Supplementary Figure 4.**
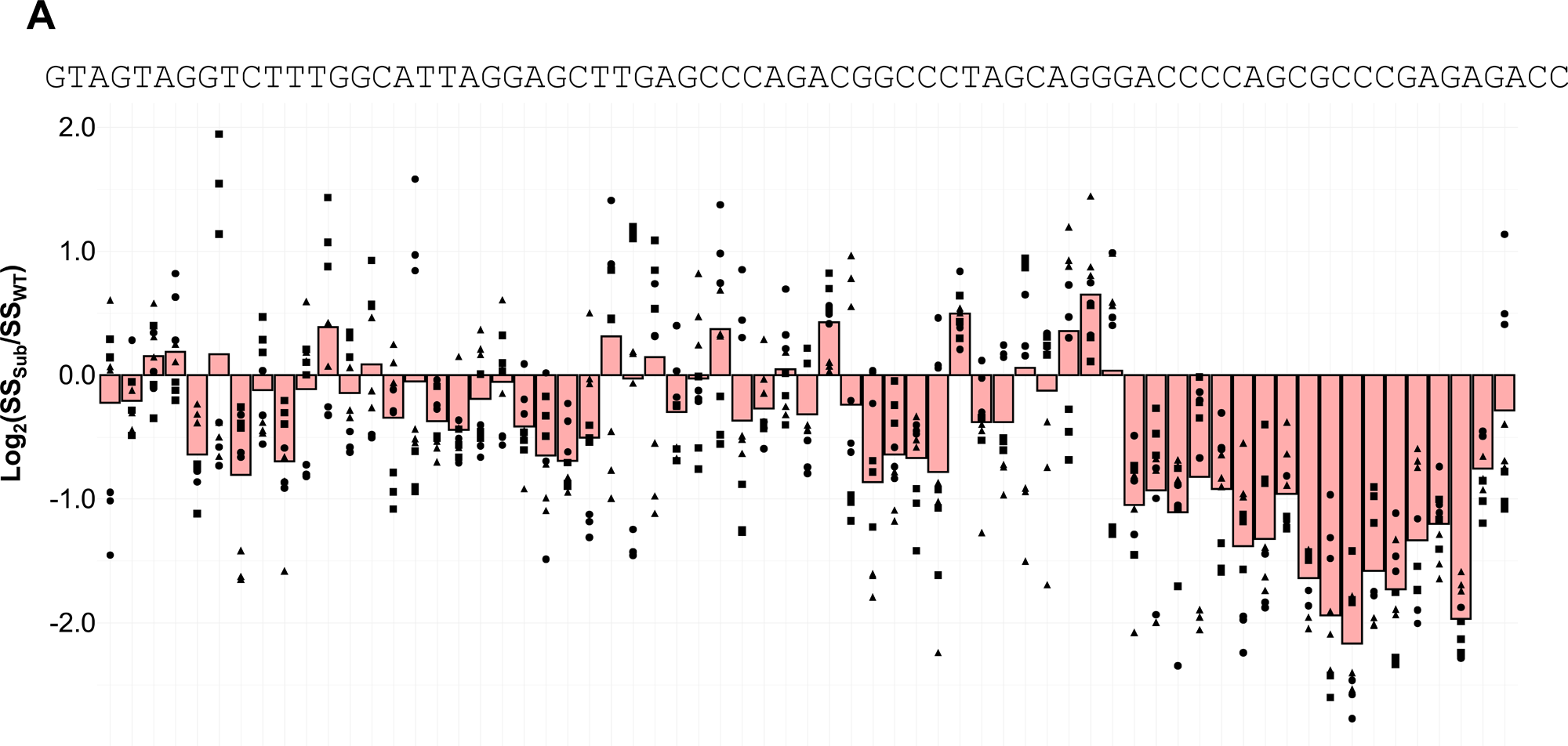
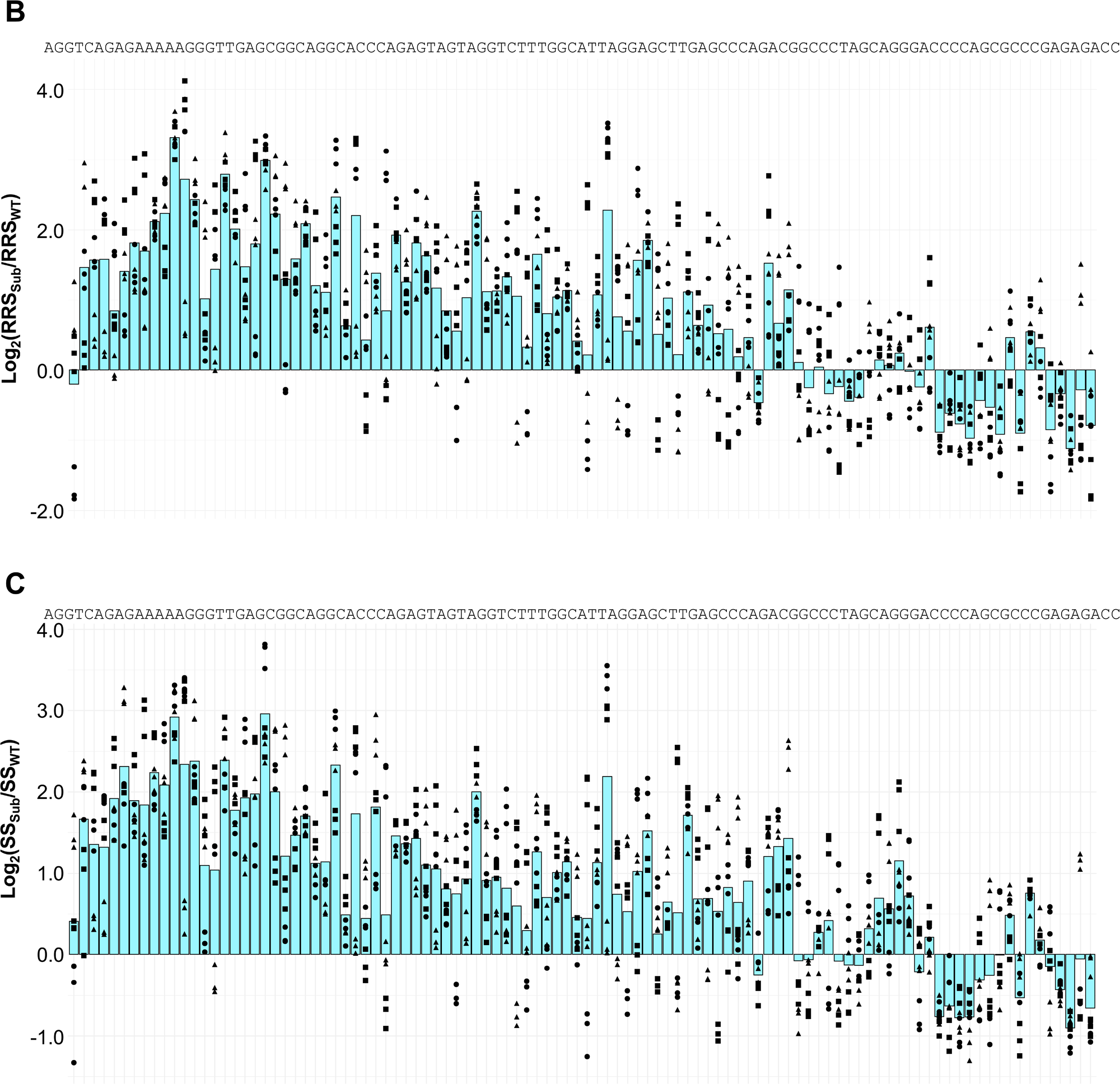
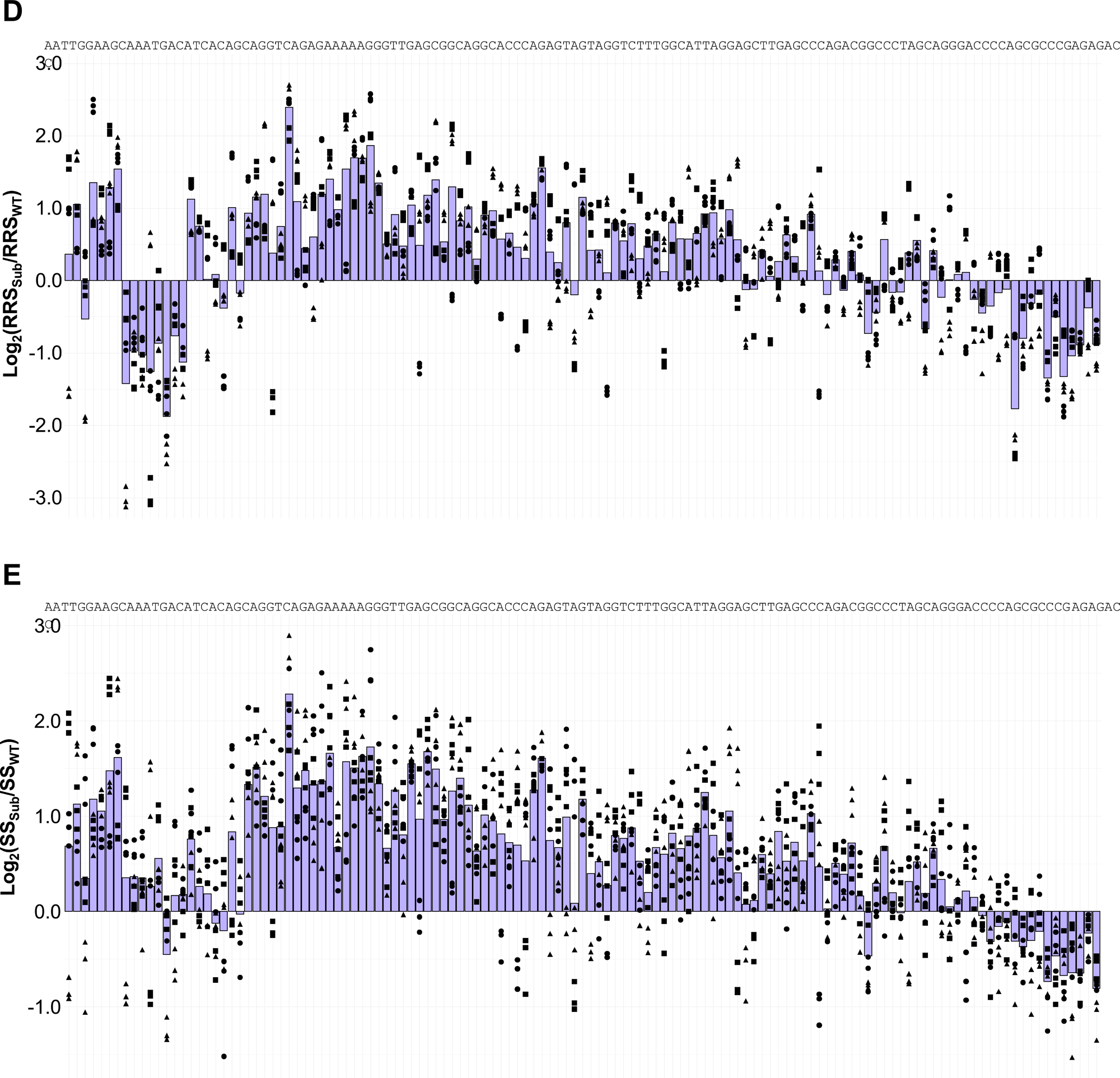
Ribosome recruitment scores (RRS) and stability scores (SS) for the CFTR-70 (**A**), CFTR-107 (**B, C**), and CFTR-132 (**D, E**) scanning substitutions. The log_2_ fold change (FC) of the RRS/SS relative to the WT was calculated for each 6-nt substitution. For each 6-nt window, three different random 6-nt substitutions were made, and all were measured by DART in three biological replicates. Bars represent the average log_2_FC averages for all replicates, and circles, squares, and triangles distinguish the three different sequences substituted at a given position. Bars are centered on the 6-nt sequence that is substituted (the WT CFTR 5’ UTR sequence shown on top).

**Supplementary Figure 5.**
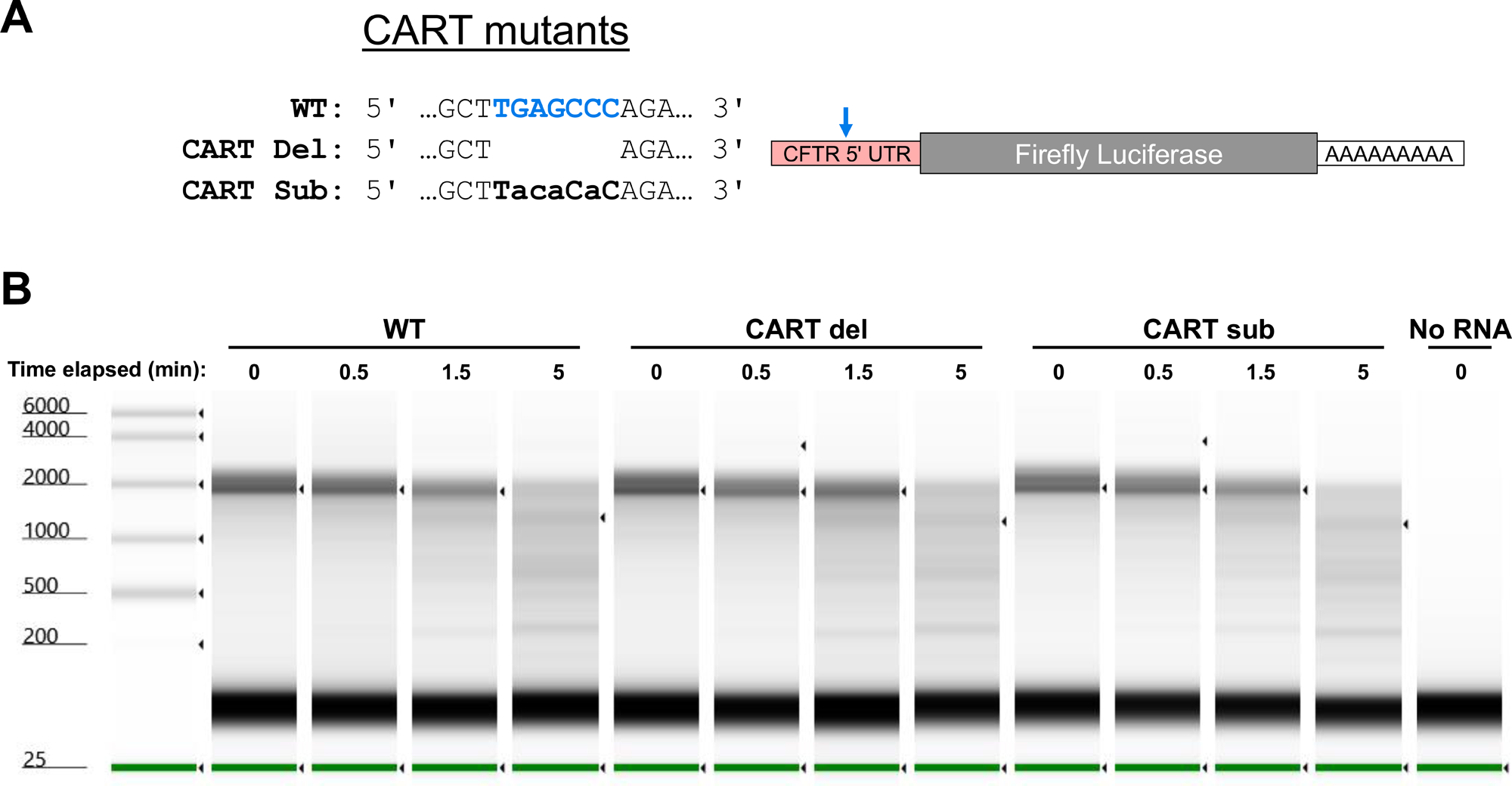
CART does not regulate inherent mRNA stability. **A.** RNA stability was measured in three luciferase reporter mRNAs; one containing the WT CFTR-70 5’ UTR and two mutants which delete (Del) or make substitutions to (Sub) the CART motif. **B.** RNA degradation was quantified by mixing mRNA with HBE cell lysate, harvesting RNA at indicated timepoints, and visualizing by RNA ScreenTape.

**Supplementary Figure 6.**
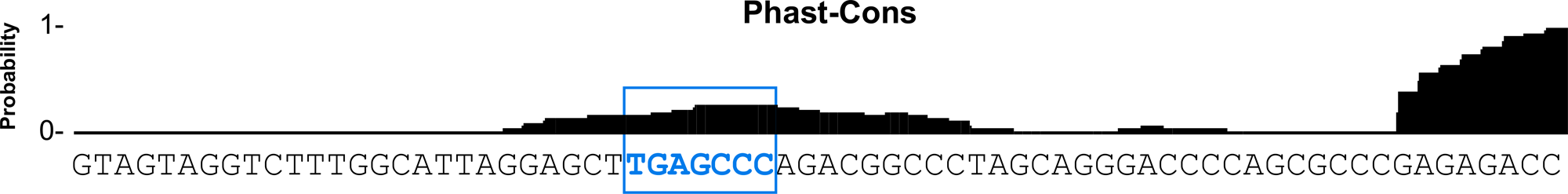
CART falls within a conserved region of the CFTR 5’ UTR. Evolutionary conservation among primates across the entire length of the CFTR-70 5’ UTR was visualized using Phast-Cons on UCSC Genome Browser.

**Supplementary Figure 7.**
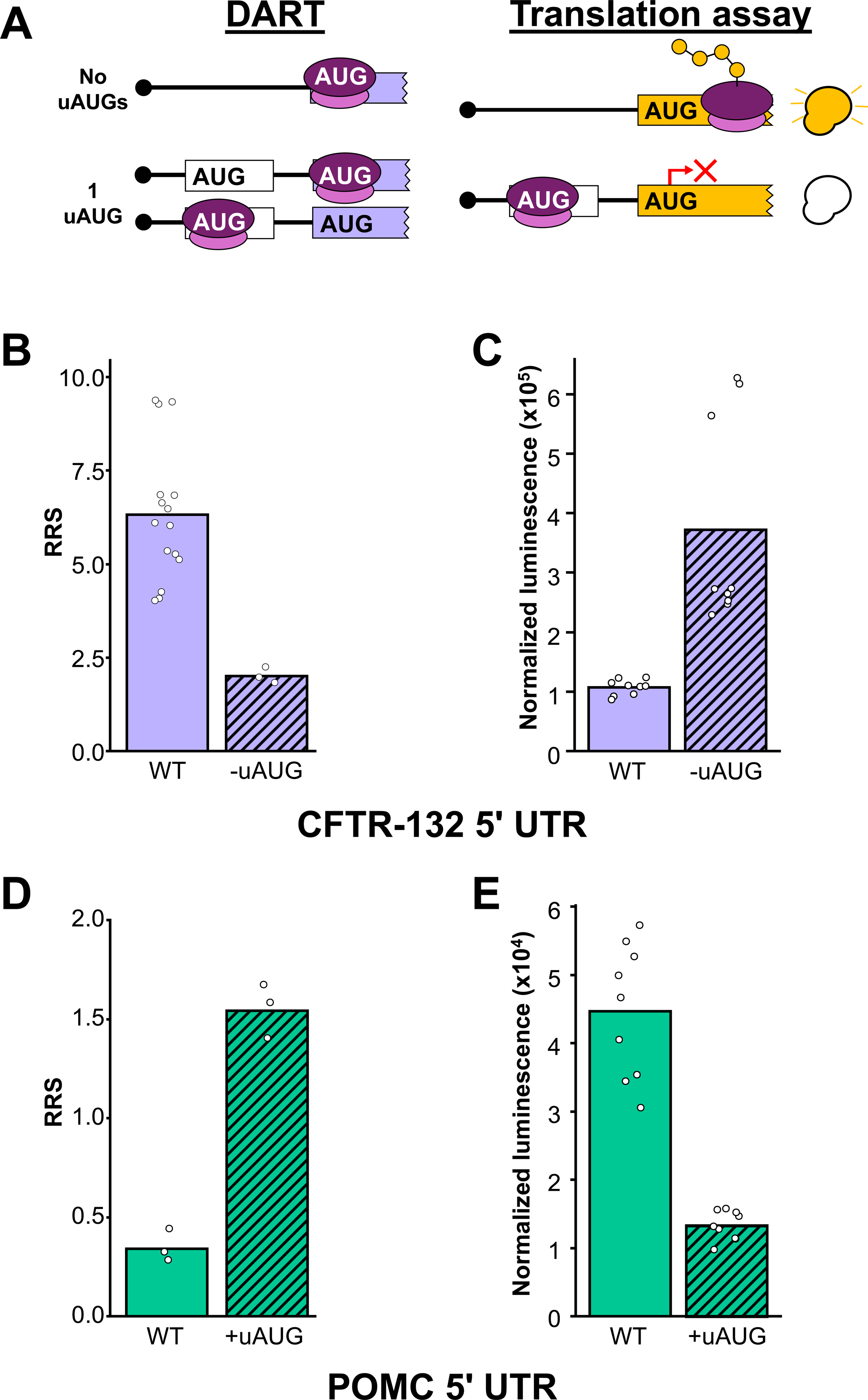
DART detects functional start sites. **A.** DART captures both uAUGs and AUGs, leading to relatively high RRS (left) even if an uAUG represses translation at the main AUG in a translation assay (right). **B.** Ribosome recruitment scores (RRS) were calculated for the wild-type CFTR-132 5’ UTR (containing an uAUG) and an uAUG→AGU mutant. Bars are averages among three biological replicates for five individually barcoded transcripts in the pool, except for the –uAUG mutant which was only included in the pool once. The assay was performed in HBE lysate. **C.** An *In vitro* translation assay was performed using luciferase reporters containing the WT CFTR-132 5’ UTR or an uAUG→AGU mutant. Bars are averages among nine biological replicates. The assay was performed using HBE lysate. **D.** As in **B** except RRS are calculated for the WT POMC 5’ UTR and a pathogenic 5’ UTR variant that forms an uAUG. Bars are averages among three biological replicates. The assay was performed in HeLa lysate. **E.** An *In vitro* translation assay was performed using luciferase reporters containing the WT POMC 5’ UTR or a pathogenic mutant which forms an uAUG. Bars are averages among nine biological replicates. The assay was performed in HeLa lysate.

**Supplementary Figure 8.**
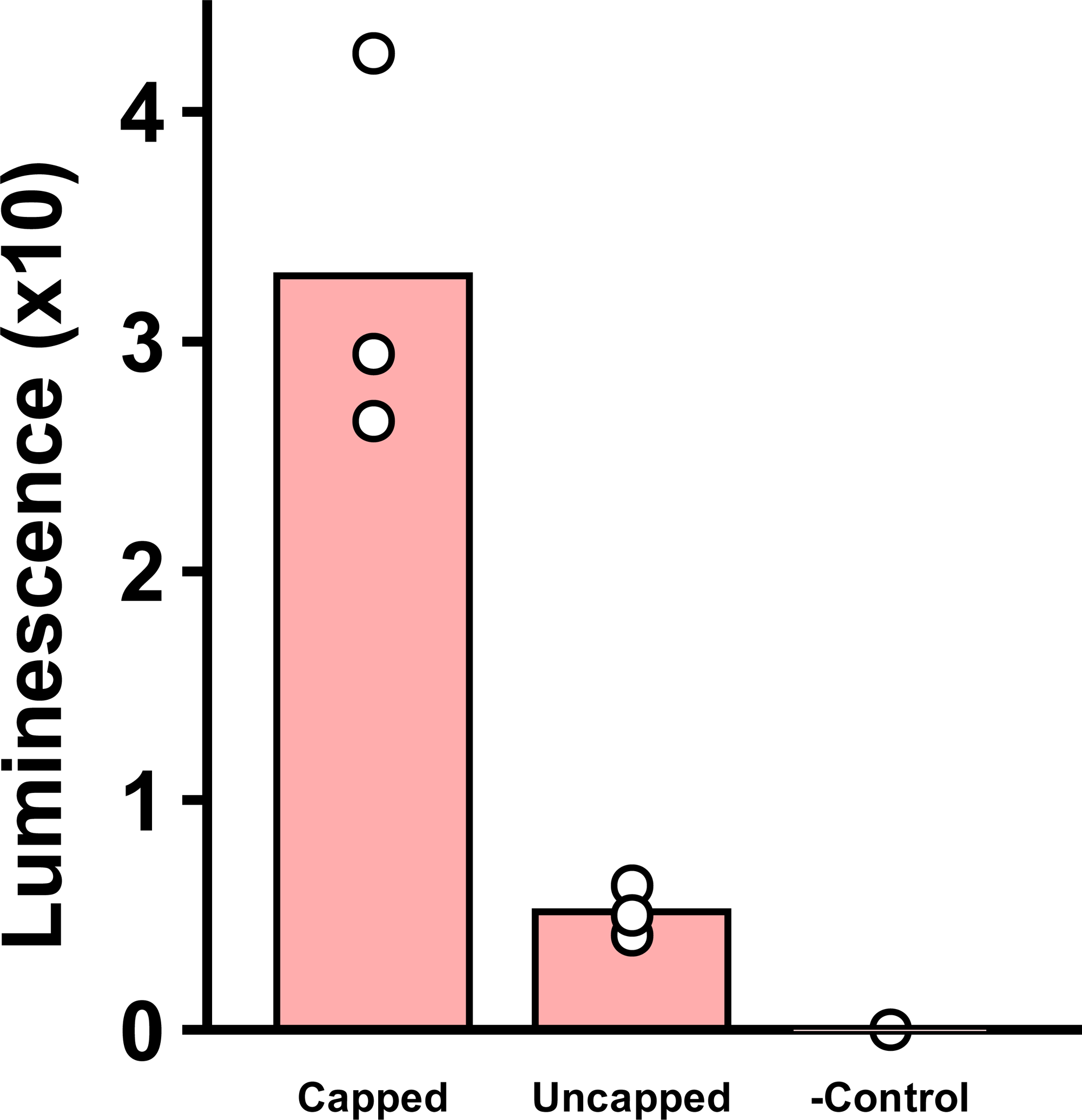
Translation of CFTR-70 is cap-dependent. An ***i****n vitro* translation assay was performed on capped and uncapped luciferase reporters containing the WT CFTR-70 5’ UTR. The negative control lacks mRNA reporter. Bars represent averages of three biological replicates. The assay was performed in HBE lysate.

## Notes

### Competing Interest Statement

The authors have declared no competing interest.

### Funding

- Cystic Fibrosis Foundation: PTAC 006048G223
- NIH Common Fund: R00GM135533

